# No observable guide-RNA-independent off-target mutation induced by prime editor

**DOI:** 10.1101/2021.04.09.439109

**Authors:** Runze Gao, Zhi-Can Fu, Xiangyang Li, Ying Wang, Jia Wei, Guangye Li, Lijie Wang, Jing Wu, Wei Xue, Xingxu Huang, Li Yang, Jia Chen

**Author notes:** Correspondence should be addressed to L.Y.; X.H.; J.C. These authors contributed equally to this work (Runze Gao, Zhi-Can Fu, Xiangyang Li, Ying Wang).

## Abstract

Prime editor (PE) has been recently developed to induce efficient and precise ontarget editing, whereas its guide RNA (gRNA)-independent off-target effects remain unknown. Here, we used whole-genome and whole-transcriptome sequencing to determine gRNA-independent off-target mutations in cells expanded from single colonies, in which PE generated precise editing at on-target sites. We found that PE triggered no observable gRNA-independent off-target mutation genome-wide or transcriptome-wide in transfected human cells, highlighting its high specificity.

## MAIN TEXT

Prime editors (PEs) that combine reverse transcriptase (RTase) with CRISPR-Cas9 system have been developed and successfully applied to induce targeted base substitutions, small deletions or insertions in mammalian cells and plants^1, 2^. By using the edit-containing genetic information encoded in the reverse transcription (RT) template of prime editing guide RNA (pegRNA), the RTase of PE can incorporate edits into target genomic DNA with high efficiency^1, 3^. Distinct to previously reported base editors (BEs), which can generate on-target C-to-T change with the combination of CRISPR-Cas9 and apolipoprotein B mRNA editing enzyme, catalytic polypeptide-like (APOBEC) cytidine deaminase, PEs are versatile to deliver different types of edits. As PE can virtually correct most of mutations associated with genetic disorders for therapeutic purposes, whether PEs induce off-target (OT) effects is of great importance for its potential clinic applications^4^. It is well studied that the engineering of Cas9 protein can greatly reduce the binding at OT sites that have sequence similarity to on-target sites^4, 5^ to reduce the gRNA-dependent OT mutations. However, whether the effector moiety of PE (RTase) induces gRNA-independent OT mutations, which were recently found to be the major OT effect for BE^6–9^, remains unknown.

At first, we sought to determine whether PE induces gRNA-independent OT mutations genome-wide. As the family of APOBEC3 cytidine deaminase has been reported to induce mutations in cellular genomic DNA^10–12^, we knocked out the endogenously expressed *APOBEC3* (*A3*) gene cluster with CRISPR-Cas9 in 293FT cells (Supplementary Fig. 1a-1d) and obtained a single-clone-derived 293FT*^A3-/-^* cell line, in order to reduce genome-wide mutation background. We next transfected the EGFP-expressing plasmid into wild-type (WT) 293FT or 293FT*^A3-/-^* cells, and 72 hours after transfection, single cells were sorted into 96-well plates. After colony expansion, the genomic DNA of single cell colonies was subjected to whole-genome sequencing (WGS) (Fig. 1a). WGS data analyzed by a <u>B</u>ase/Prime <u>e</u>ditor <u>i</u>nduced <u>D</u>NA <u>o</u>ff-target site identification <u>u</u>nified toolkit (BEIDOU, https://github.com/YangLab/BEIDOU) showed that the knockout of *A3* cluster significantly reduced the number of genome-wide base substitutions (Supplementary Fig. 2a, from ∼4,900 to ∼1,600, *P* = 6 × 10^-5^) and insertions or deletions (indels, Supplementary Fig. 2b, from ∼400 to ∼20, *P* = 2 × 10^-8^). In the subsequent study, we then used the 293FT*^A3-/-^* cell line with a low mutation background to evaluate the gRNA-independent OT effects induced by PEs genome-wide, with EGFP as the negative control and a previously reported BE (hA3A-BE3)^13^ as the positive control for detecting gRNA-independent OT mutations.

**Fig. 1.**
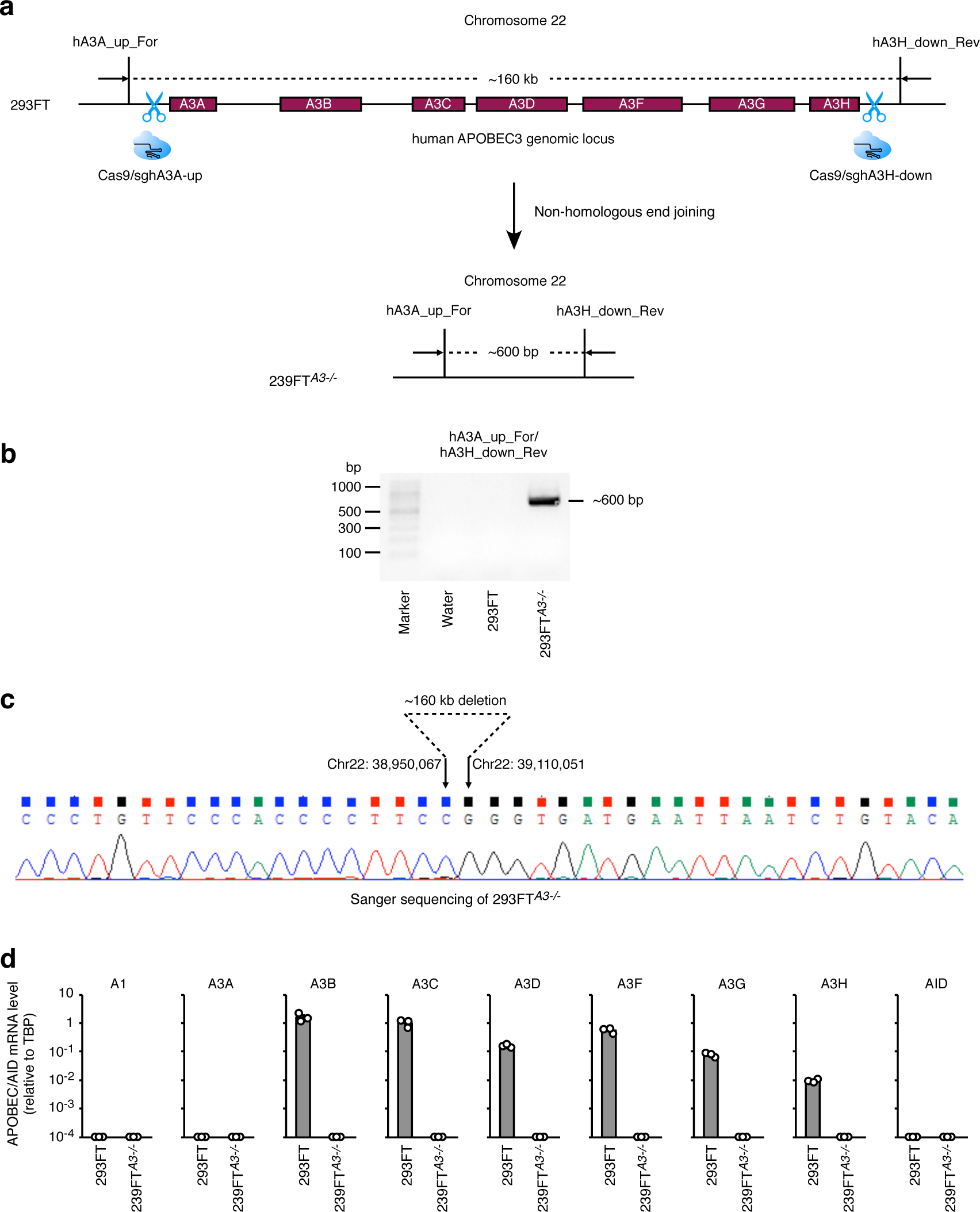
PE3 induced no observable genome-wide OT mutation when generating base substitutions. (**a**) Schematic diagrams illustrate the procedure to determine genome-wide OT mutations in edited single cell colonies. Briefly, 293FT*^A3-/-^* cells were transfected with genome editors and after transfection, single cells were sorted into 96-well plates by cell sorter. After colony expansion, the genomes derived from single cell colonies were extracted for Sanger sequencing and then the ones with bi-allelic editing were further subjected to WGS. (**b**) Schematic diagrams illustrate *RNF2*, *FANCF* and *SEC61B* target sites, the spacer sequences of gRNAs and the primer-binding sites and reverse transcription (RT) templates of pegRNAs. The editing sites of PE3 and hA3ABE3 are in red and green respectively. (**c**) C-to-T editing frequencies (count of reads with C-to-T editing at the target base/count of reads covering the target base) induced by the indicated genome editors at target sites. Means ± s.d. were from three independent experiments. (**d**) Numbers of genome-wide base substitutions induced by indicated genome editors in single cell colonies with bi-allelic edits at on-target sites shown in Supplementary Fig. 4. (**e**) Numbers of each type of base substitution induced by indicated genome editors shown in (**d**). (**f**) Numbers of genome-wide indels induced by indicated genome editors in single cell colonies shown in Supplementary Fig. 4. (**d**) - (**f**) Means ± s.d. were from four or six (EGFP) independent colonies. *P* value, one-tailed Student’s *t* test.

We first compared the on-target editing efficiencies of PE3 to hA3A-BE3 with amplicon sequencing in a bulk transfection setting. 293FT*^A3-/-^* cells were transfected with three plasmids expressing PE3, pegRNA (Fig. 1b) and the optimized nicking gRNA^1^ (Supplementary Fig. 3a), two plasmids expressing hA3A-BE3 and gRNA, two plasmids expressing Cas9 and gRNA, or one plasmid expressing EGFP. 72 hours after transfection, a portion of transfected cells was lysed to extract genomic DNA for examining on-target editing frequencies. PE3 generated C-to-T substitutions at *RNF2*, *FANCF* and *SEC61B* target sites, with editing efficiencies similar to those by hA3A-BE3 (Fig. 1c). Consistent with previous studies showing that both PE3 and hA3ABE3 triggered indels^1, 13^, we found that both editors induced indels at these on-target sites, but fewer than those by Cas9 (Supplementary Fig. 3b).

Another portion of transfected cells was sorted to single cells and expanded in 96-well plates. After culturing, the single cell colonies with Sanger-sequencing confirmed biallelic edits (100% editing frequency) at on-target sites (Supplementary Fig. 4a-4c) were subjected to WGS with a depth of at least 12× to identify OT mutations and analyzed by the BEIDOU toolkit on a genome-wide scale (Supplementary Fig. 5a-5d). We found the numbers of base substitutions in PE3-treated (Fig. 1d, *P* = 0.13, 0.08 and 0.02, for *RNF2*, *FANCF* and *SEC61B*, respectively) and Cas9-treated single cell colonies (Fig. 1d, *P* = 0.24, 0.02 and 0.14, for *RNF2*, *FANCF* and *SEC61B*, respectively) were similar to or slightly smaller than the ones in EGFP-treated single cell colonies, suggesting that PE3 and Cas9 induced no observable genome-wide OT base substitution. In contrast, hA3A-BE3 induced more base substitutions than EGFP control (Fig. 1d, *P* = 0.004, 0.13 and 4 × 10^-5^, for *RNF2*, *FANCF* and *SEC61B*, respectively), which is in line with previous reports that BE3 induced substantial genome-wide OT mutations in mouse embryos and plants^7, 9^. We further analyzed the type of base substitutions and found that significantly more C-to-T or G-to-A substitutions than other subtypes of base substitutions (Fig. 1e and Supplementary Fig. 6a, *P* < 0.008) were induced by hA3A-BE3, while C-to-T or G-to-A substitutions were not significantly induced by other genome editors (Fig. 1e and Supplementary Fig. 6b). We also analyzed sequence contexts of base substitutions induced by hA3ABE3 and found that the WGS-identified OT mutation sites have little sequence similarity to on-target sites or predicted gRNA-dependent OT sites (Supplementary Fig. 7a-7f). This finding is consistent with previous studies^7, 9, 14, 15^ and indicates that these OT mutations are gRNA-independent and randomly induced by the cytidine deaminase moiety of hA3A-BE3^16, 17^. Moreover, we used Sanger sequencing to confirm the WGS-identified gRNA-independent OT mutations at selected loci that are associated with human diseases and found ∼50% mutation frequencies at these pathogenic sites in single cell colonies (Supplementary Fig. 8a and 8b). Other than base substitutions, we also analyzed whether PE3 and hA3A-BE3 induced gRNA-independent OT indels genome-wide when these two editors were used to generate on-target base substitutions. Both of them only manifested background numbers of indels, comparing to Cas9 or EGFP control (Fig. 1f and Supplementary Fig. 9a-9d). These results together demonstrated that PE3 induced no observable gRNA-independent OT base substitution or indel in its application of introducing single base changes.

An advantage of applying PE3 for genome editing is to introduce small deletions or insertions at targeted sites^1, 2^. Thus, we further set to evaluate gRNA-independent OT effects in the application of PE3 for generating designed deletions or insertions. With the same strategy shown in Fig. 1a, we transfected 293FT*^A3-/-^* cells with PE3, Cas9 and EGFP. With three pairs of pegRNA and nicking gRNA with optimal editing efficiencies^1^ (Supplementary Fig. 10a), PE3 was used to generate 3-bp deletions at *EMX1*, *HEK1* and *LSP1* sites (Fig. 2a) and deep sequencing results showed that PE3 yielded ∼40%-70% intended deletion frequencies in a bulk transfection setting (Fig. 2b). Next, the whole genomes of edited single cell colonies with intended 3-bp deletions (Supplementary Fig. 11a-11c) were sequenced and then analyzed by the BEIDOU toolkit (Supplementary Fig. 12a and 12b). WGS results showed that PE3 induced similar or slightly fewer OT indels, comparing to Cas9 (Fig. 2c, *P* = 0.32, 0.15 and 0.06 for *EMX1*, *HEK1* and *LSP1*, respectively) or EGFP control (Fig. 2c, *P* = 0.14, 0.11 and 0.03 for *EMX1*, *HEK1* and *LSP1*, respectively). Meanwhile, the numbers of base substitutions induced by PE3 were similar to or slightly smaller than Cas9 (Fig. 2d, *P* = 0.21, 0.23 and 0.001 for *EMX1*, *HEK1* and *LSP1*, respectively) and EGFP control (Fig. 2d, *P* = 0.07, 0.04 and 0.02 for *EMX1*, *HEK1* and *LSP1*, respectively), consistent to the results shown in Fig. 1. We further confirmed that these designed 3-bp deletions were not induced at OT sites on a genome-wide scale (Supplementary Fig. 13a-13c).

**Fig. 2.**
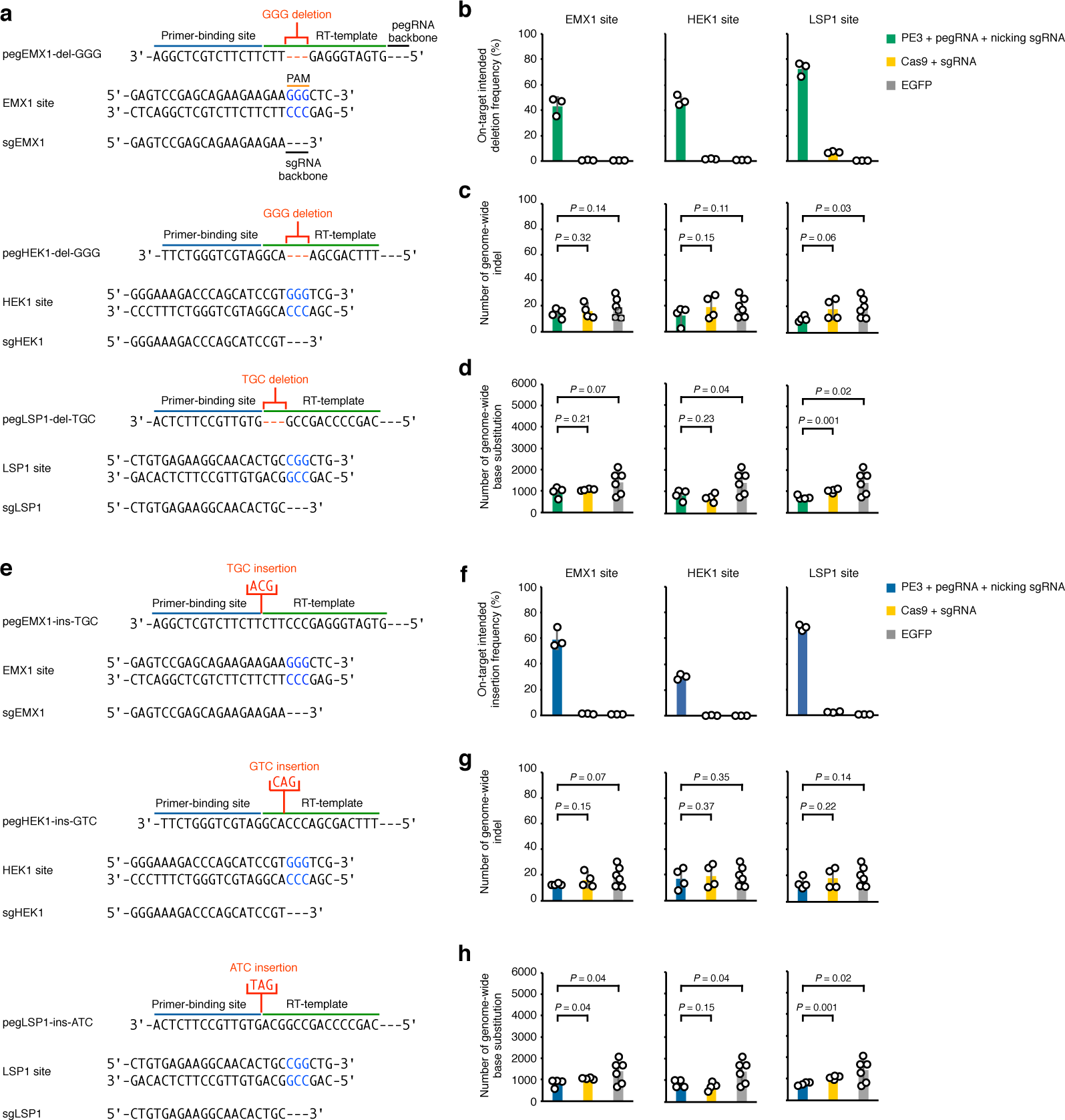
PE3 induced no observable genome-wide OT mutation when generating small deletions and insertions. (**a**) Schematic diagrams illustrate *EMX1*, *HEK1* and *LSP1* target sites, the spacer sequences of gRNAs and the primer-binding sites and RT templates of pegRNAs for targeted deletions. The designed deletions in pegRNA are in red. (**b**) Targeted deletion frequencies induced by the indicated genome editors at on-target sites. (**c**), (**d**) Numbers of genome-wide indels (**c**) and base substitutions (**d**) induced by indicated genome editors in single cell colonies shown in Supplementary Fig. 11. (**e**) Schematic diagrams illustrate *EMX1*, *HEK1* and *LSP1* target sites, the spacer sequences of gRNAs and the primer-binding sites and RT templates of pegRNAs for targeted insertions. The designed insertions in pegRNA are in red. (**f**) Targeted insertion frequencies induced by the indicated genome editors at on-target sites. (**g**), (**h**) Numbers of genome-wide indels (**g**) and base substitutions (**h**) induced by indicated genome editors in single cell colonies shown in Supplementary Fig. 15. The data for Cas9 and EGFP in (**f**)-(**h**) are same as the ones in (**b**)-(**d**). (**b**) and (**f**) Means ± s.d. were from three independent experiments. (**c**), (**d**), (**g**) and (**h**) Means ± s.d. were from four or six (EGFP) independent colonies. *P* value, one-tailed Student’s *t* test.

**Fig. 3.**
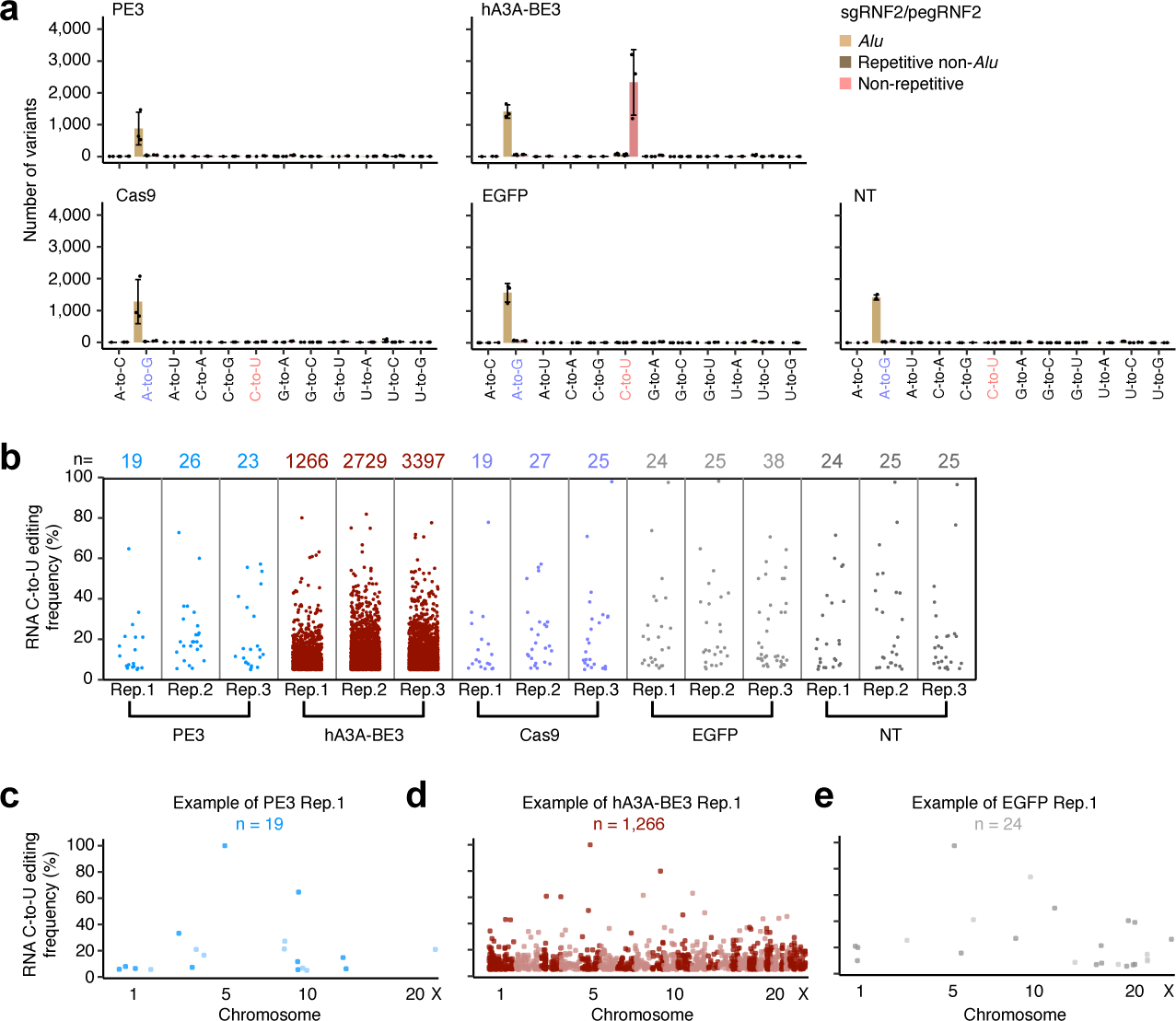
PE3 induced no observable transcriptome-wide OT mutation. (**a**) Histogram to show numbers of all 12 types of RNA editing in different regions from cells treated with PE3, hA3A-BE3, Cas9, EGFP or left non-transfected (NT). RNA editing was analyzed and visualized by the RADAR pipeline. Means ± s.d. were from three independent experiments. (**b**) Manhattan plot of RNA off-target editing (C-to-U) frequency shown in (**a**). (**c**, **d, e**) The RNA off-target editing frequencies and sites induced

Finally, we used PE3 and three pairs of pegRNA and nicking gRNA with optimal editing efficiencies^1^ (Supplementary Fig. 14a) to generate 3-bp insertions at *EMX1*, *HEK1* and *LSP1* genomic sites (Fig. 2e). Deep sequencing showed that PE3 induced ∼30%-70% intended insertion frequencies (Fig. 2f) at the on-target sites. WGS of the PE3-edited single cell colonies (Supplementary Fig. 15a-15c) displayed that PE3 induced similar or fewer indels (Fig. 2g and Supplementary Fig. 16a) and base substitutions (Fig. 2h and Supplementary Fig. 16b), comparing to Cas9 or EGFP control. Meanwhile, the designed 3-bp insertions were not found at OT sites on a genome-wide scale (Supplementary Fig. 17a-17c).

Furthermore, we transfected different amounts of plasmids to express PE, pegRNA and nicking gRNA or transfected same amounts of plasmids but for different exposure time periods. In all tested conditions, PE3 manifested no more genome-wide OT mutations than the EGFP control (Supplementary Fig. 18a-18c). We also examined whether PE3 copied pegRNA scaffold into genome and found that no pegRNA sequence was inserted into the genomic DNA (Supplementary Fig. 19a and 19b). As the effector moiety of PE3 is an RTase, we therefore determined whether PE3 affected telomere integrity in PE3-treated single cell colonies and discovered that the length and sequence of telomeric region were barely affected by PE3 (Supplementary Fig. 20a-20d). After examing the gRNA-independent OT effects genome-wide, we also determined the gRNA-dependent OT mutations by PE3 in WT 293FT cells. No obvious base substitution (Supplementary Fig. 21a and 21b) or indel (Supplementary Fig. 22a and 22b) was induced by PE3 at gRNA-dependent OT sites^18, 19^. Nevertheless, it is worthwhile noting that although PE3 induced relatively lower levels of unintended indels at on-target sites than Cas9, the average unintended indel frequencies induced by PE3 at on-target sites were ∼10% (Supplementary Fig. 3b, 10b and 14b). It is likely due to the requirement of both pegRNA and nicking gRNA in the PE3 system, which may cleave opposite DNA strands near on-target site to generate a DSB, and the repair of resulted DSB could then lead to small indels or possibly, large genomic deletions^20^.

In addition to gRNA-independent OT effects on genomic DNA, gRNA-independent OT effects of genome editors on transcriptomic RNA have been recently reported as well^6, 8^. Thus, we performed whole-transcriptome sequencing and used the RADAR pipeline^21^ to detect whether PE3 induced gRNA-independent OT mutations in transcriptomic RNA in WT 293FT cells, with EGFP and Cas9 as negative controls and hA3A-BE3 as a positive control. Whole-transcriptome sequencing results demonstrated that comparing to EGFP or Cas9, PE3 induced similar mutations at the transcriptomic RNA level (Fig. 3a, 3b, 3c and 3e). In contrast, hA3A-BE3 induced much more mutations in transcriptomic RNA than EGFP, Cas9 and PE3 (Fig. 3a, 3b and 3d), and most of mutations were C-to-U substitutions (Fig. 3a), which were catalyzed by the hA3A deaminase moiety of hA3A-BE3 in a gRNA-independent manner. Furthermore, we also examined whether PE treatment affects gene expression and found that expression patterns in PE-treated and EGFP-treated cells were similar, suggesting that PE did not affect gene expression (Supplementary Fig. 23a and 23b).

Collectively, we evaluated gRNA-independent OT mutations of PE3 by performing whole-genome and whole-transcriptome sequencing and showed that PE3 induced no observable OT mutation in a gRNA-independent manner, when generating three types of targeted edits, *i.e.*, base substitution, small deletion and insertion. Together with previous studies showing the versatility and high efficiency of PE’s on-target editing^1^, our results highlighted the high editing specificity of PE system.

## METHODS

### Plasmid construction

Oligonucleotides hRNF2_FOR/hRNF2_REV were annealed and ligated into BsaI linearized pGL3-U6-gRNA-PGK-EGFP to generate the vector psgRNF2 for the expression of sgRNF2. Oligonucleotides +41_nicking-hRNF2 _FOR/+41_nickinghRNF2 _REV were annealed and ligated into BsaI linearized pGL3-U6-gRNA-PGKpuromysin to generate the vector pnsgRNF2 for the expression of +41_nicking sgRNF2. Other gRNA and nicking gRNA expression vectors were constructed by a similar strategy, respectively.

The primer sets (pegRNF2_F/pegRNF2_R) were used to amplify the fragment scaffold-pegRNF2 with the template pU6-pegRNA-GG-Vector^1^.Then, the amplified fragment scaffold-pegRNF2 was cloned into BsaI and EcoRI linearized pU6pegRNA-GG-Vector to generate the vector pU6-pegRNF2. Other pegRNA expression vectors were constructed by a similar strategy, respectively.

The sequences of the oligos used for plasmid construction are listed in Supplementary Table 1.

### Establishment of 293FT*^A3-/-^* cells

To establish the *APOBEC3*-knockout (293FT*^A3-/-^*) cell line, 293FT cells were seeded into a 60-mm plate at a density of 4 × 10^5^ per well and cultured for 24 hours. The cells were then transfected with a plasmid expressing Cas9 nuclease and a plasmid expressing two gRNAs targeting the upstream genomic region of *APOBEC3A* and the downstream genomic region of *APOBEC3H* (sghA3A-up and sghA3H-down, Supplementary Fig. 1a), with a puromycin-resistant selection gene. After 48 hours, 10 μg/ml puromycin was added into the media to enrich the transfected cells and after two weeks of enrichment, single cell colonies were sorted into 96-well plates without puromycin for colony expansion. The non-homologous end joining of two DNA ends generated by Cas9 and sghA3A-up/sghA3H-down can remove the whole gene cluster of *APOBEC3* from the genome of 293FT cells and the single cell colony containing successful knockout of *APOBEC3* cluster was validated by genomic DNA PCR with the primers flanking the Cas9 cleavage sites (Supplementary Fig. 1b and 1c). RTqPCR^22^ further confirmed that the expression of all *APOBEC3* genes was not detected in 293FT*^A3-/-^* cells (Supplementary Fig. 1d).

### Cell culture and transfection for on-target and gRNA-dependent off-target deep-sequencing

293FT from ATCC or 293FT*^A3-/-^* cells were maintained in DMEM (10566, Gibco/Thermo Fisher Scientific) + 10% FBS (16000-044, Gibco/Thermo Fisher Scientific) and regularly tested to exclude mycoplasma contamination.

Before transfection, cells were seeded in a 24-well plate at a density of 1× 10^5^ per well. For base editing with hA3A-BE3 and gene editing with Cas9, the cells were transfected with 250 μl serum-free Opti-MEM that contained 2.52 μl LIPOFECTAMINE LTX (Life, Invitrogen), 0.84 μl LIPOFECTAMINE plus (Life, Invitrogen), 0.5 μg pCMV-hA3A-BE3 (or pCMV-spCas9) expression vector and 0.34 μg gRNA expression vector. For prime editing with PE3, the cells were transfected with 250 μl serum-free Opti-MEM that contained 3.9 μl LIPOFECTAMINE LTX, 1.3 μl LIPOFECTAMINE plus, 0.9 μg PE3 expression vector, 0.3 μg pegRNA expression vector and 0.1 μg nicking gRNA expression vector. For EGFP expression, the cells were transfected with 250 μl serum-free Opti-MEM that contained 1.5 μl LIPOFECTAMINE LTX (Life, Invitrogen), 0.5 μl LIPOFECTAMINE plus (Life, Invitrogen), 0.5 μg pCMV-EGFP expression vector. 72 hours after transfection, transfected cells in the first 10% of the fluorescence intensity were sorted by BD FACSAria III, from which the genomic DNA was extracted with QuickExtract^TM^ DNA Extraction Solution (QE09050, Epicentre) for subsequent sequencing analysis.

### Isolation and expansion of edited single cell colonies for whole-genome sequencing

293FT*^A3-/-^* cells expanded from a single cell colony with successful *A3* knockout were maintained in DMEM (10566, Gibco/Thermo Fisher Scientific) + 10% FBS (16000-044, Gibco/Thermo Fisher Scientific) and regularly tested to exclude mycoplasma contamination. The single cell colony-derived 293FT*^A3-/-^* cells were transfected with genome editors (*e.g.*, PE3, hA3A-BE3 and Cas9) or EGFP-expressing plasmids and 72 hours after transfection, single cells were sorted into 96-well plates by BD FACSAria III. After 18-day colony expansion, the genomic DNA derived from transfected single cell colonies was extracted with QuickExtract^TM^ DNA Extraction Solution (QE09050, Epicentre) for Sanger sequencing and the genomic DNA with biallelic editing was further subjected to whole-genome sequencing. On average, one bi-allelic edited colony can be obtained from ∼6-8 colonies when the editing efficiency of bulk setting is ∼30-50%.

### DNA library preparation and sequencing

Target genomic sequences were PCR amplified by high-fidelity DNA polymerase PrimeSTAR HS (Clonetech) with primer sets flanking examined gRNA target sites. The gRNA target sequences and PCR primer sequences are listed in Supplementary Table 2. Indexed DNA libraries were prepared by using the NEBNext Ultra II FS DNA Library Prep Kit for Illumina. After quantitated with Qubit High-Sensitivity DNA kit (Invitrogen), PCR products with different tags were pooled together for deep sequencing by using the Illumina Hiseq X Ten (2×150) at CAS-MPG Partner Institute for Computational Biology Omics Core, Shanghai, China. Raw read qualities were evaluated by FastQC (v0.11.8, http://www.bioinformatics.babraham.ac.uk/projects/fastqc/, parameters: default). For paired-end sequencing, only R1 reads were used. Adaptor sequences and read sequences with Phred quality score lower than 30 were trimmed. Trimmed reads were then mapped with the BWA-MEM algorithm (BWA v0.7.17) to target sequences. After piled up with Samtools (v1.9), base substitutions and indel frequencies at ontarget sites were calculated according to previously published literature^13, 23^.

### Base substitution frequency calculation

Base substitution of every position at the target sites of examined gRNAs and pegRNAs was piled up with at least 1000 independent reads. Base substitution frequencies were calculated as: (count of reads with substitution at the target base)/(count of reads covering the target base). Counts of reads for each base at examined target sites and gRNA-dependent OT sites are listed in Supplementary Table 3 and 5, respectively.

### Intended indel frequency calculation

Intended indel frequencies were calculated as: (count of reads with only intended indel at the target site)/(count of total reads covering the target site). These counts are listed in Supplementary Table 4.

### Unintended indel frequency calculation

Unintended indel frequencies for base substitution were estimated among reads aligned in the region spanning from upstream 8 nucleotides to the target site (or gRNA-dependent OT site) to downstream 19 nucleotides to PAM site (50 bp). Unintended indel frequencies for base substitution were calculated as: (count of reads containing at least one unintended inserted and/or deleted nucleotide)/(count of total reads aligned in the estimated region). While unintended indel frequencies for targeted insertion/deletion were estimated among reads aligned at the target site. Unintended indel frequencies for targeted insertion/deletion were calculated as: (count of reads containing unintended indels)/(count of total reads covering the target site). The counts for on-target and gRNA-dependent OT sites are listed in Supplementary Tables 4 and 5, respectively.

### Whole-genome sequencing and data analysis

Genomic DNA was extracted from transfected 293FT*^A3-/-^* single cell colonies by using cell DNA isolation kit FastPure^®^ (DC102-01, Vazyme). Indexed DNA libraries were prepared by using NEBNext Ultra II FS DNA Library Prep Kit for Illumina. A total of 12 Tb WGS data were obtained by using Illumina Hiseq X Ten (2×150) at CASMPG Partner Institute for Computational Biology Omics Core, Shanghai, China. The average coverage of sequencing data generated for each transfected 293FT^*A3-/-*^ single cell colony sample was 14×, with a minimum depth at 12×. These WGS datasets were individually analyzed with a <u>B</u>ase/Prime <u>e</u>ditor <u>i</u>nduced <u>D</u>NA <u>o</u>ff-target site identification <u>u</u>nified toolkit (BEIDOU, https://github.com/YangLab/BEIDOU) to call high-confident base substitution or indel events that could be identified by all three different callers, GATK^24^, Lofreq^25^ and Strelka2^26^.

Briefly, to reduce the impact of varying sequence depth among samples, 120M reads were randomly sampled by Seqtk (v1.3, https://github.com/lh3/seqtk, parameters: sample -s100 120000000) from raw data for further analyses. After quality control by FastQC (parameters: default), WGS DNA-seq reads were trimmed by Trimmomatic (v0.38, parameters: ILLUMINACLIP:TruSeq3-PE-2.fa: 2:30:10 LEADING:3 TRAILING:3 SLIDINGWINDOW:4:15 MINLEN:36)^27^ to remove low quality read sequence. BWA-MEM algorithm (v0.7.17, parameters: default) was used to map clean reads to the human reference genome (hg38). Samtools (v1.9, parameters: -bh F 4 -q 30) was used to select reads with mapping quality score ≥ 30 and convert SAM files to sorted BAM files. After marking duplicate reads by Picard (v2.21.2, parameters: REMOVE_DUPLICATES=false) in the BAM file, GATK (v4.1.3.0) was employed to correct systematic bias by a two-stage process (BaseRecalibrator and ApplyBQSR, parameters: default).

Single nucleotide variations of OT mutations were individually computed by the BEIDOU toolkit with three algorithms GATK, Lofreq (v2.1.3.1, parameters: default) and Strelka2 (v2.9.10, parameters: default) with workflows for the germline variant calling. Genome-wide indels were also detected by the BEIDOU toolkit with GATK, Strelka2 (parameters: default) and Scalpel (v0.5.4, parameters: --single --window 600)^28^. For GATK, genome-wide *de novo* variants were determined by three GATK commands, HaplotypeCaller (parameters: default), VariantRecalibrator (parameters: “--resource:hapmap,known=false,training=true,truth=true,prior=15.0 hapmap_3.3.hg38.vcf.gz -resource:omni,known=false,training=true,truth=false,prior=12.0 1000G_omni2.5.hg38.vcf.gz -resource:1000G,known=false,training=true,truth=false,prior=10.0 1000G_phase1.snps.high_confidence.hg38.vcf.gz -resource:dbsnp,known=true,training=false,truth=false,prior=2.0 dbsnp_146.hg38.vcf.gz -an QD -an MQ -an MQRankSum -an ReadPosRankSum -an FS -an SOR -an DP --max-gaussians 4” for SNVs; “resource:mills,known=true,training=true,truth=true,prior=12.0 Mills_and_1000G_gold_standard.indels.hg38.vcf.gz -an QD -an MQRankSum -an ReadPosRankSum -an FS -an SOR -an DP --max-gaussians 4 -mode INDEL” for indels) and ApplyVQSR (parameters: “-mode SNP -ts-filter-level 95” for SNVs; “mode INDEL -ts-filter-level 95” for indels). VCF files used for VariantRecalibrator were downloaded from https://ftp.ncbi.nih.gov/snp/ and https://console.cloud.google.com/storage/browser/genomics-publicdata/resources/broad/hg38/v0. Of note, overlaps of three algorithms of SNVs/indels were considered as reliable variants by the BEIDOU toolkit. To further obtain *de novo* SNVs/indels, we filtered out the background variants, including: (1) SNVs/indels in non-transfected cells of this study and dbSNP (v151, http://www.ncbi.nlm.nih.gov/SNP/) database; (2) SNVs/indels with allele frequencies less than 10% or depth less than 10 reads; (3) SNVs/indels overlapped with the UCSC repeat regions. Analyses were only focused on SNVs/indels from canonical (chr 1–22, X, Y and M) chromosomes. Genome-wide base substitutions and indels are listed in Supplementary Table 6.

### Predication of gRNA-dependent OT site

Potential gRNA-dependent OT sites were predicted by Cas-OFFinder^19^, allowing up to 5 mismatches. OT sites identified by WGS shown in Supplementary Fig. 7 were randomly selected from genome-wide base substitutions.

### Telomere length calculation and variant calling from WGS data

Telseq^29^ (parameters: -k 2) was used to calculate the telomere lengths. BAM files containing mapped WGS DNA-seq reads processing from the BEIDOU toolkit were as the input of Telseq. Telomeric repeat variants, the sequence fragments within telomeric reads that differ from the canonical telomeric repeat pattern (TTAGGG in human), were identified by Computel^30^ (v1.2, parameters: default) with trimmed WGS fastq files as input. Telomere length and the numbers of telomeric repeat variants are listed in Supplementary Table 7.

### RNA extraction and whole-transcriptome sequencing and data analysis

After 48 hours of transfection with genome editors (*e.g.*, PE3, hA3A-BE3 and Cas9) or EGFP-expressing plasmids, transfected cells in the first 10% of the fluorescence intensity were sorted by BD FACSAria III. Total RNAs of sorted cells were extracted by using the RNeasy Mini Kit (QIAGEN #74104) for whole-transcriptome sequencing. RNA-seq libraries were prepared using Illumina TruSeq Stranded Total RNA LT Sample Prep Kit. Size-selected libraries were subjected to deep sequencing with Illumina Hiseq X Ten (2×150). Raw read qualities were evaluated by FastQC (http://www.bioinformatics.babraham.ac.uk/projects/fastqc/). RNA editing sites were called by the published RADAR pipeline^21^. Gene expression was determined by FPKM (Fragments Per Kilobase of transcript per Million mapped reads) with featureCounts (v1.6.3, --fraction -O -t exon -g gene_id). Transcriptome-wide mutations are listed in Supplementary Table 8.

### Data Availability

Deep sequencing data and WGS data can be accessed in the NCBI Gene Expression Omnibus (accession no. xxxx) and the National Omics Data Encyclopedia (accession no: OEPxxxxxx. Of note, the accession code will be available before publication).

### Code Availability

The BEIDOU tookit that calls high-confident base substitution or indel events from WGS data is available at https://github.com/YangLab/BEIDOU. The custom Python, Perl and Shell scripts for base substitution and indel frequencies calculation will be available upon request.

### Statistical analysis

All statistical analyses were performed with R package 3.6.2 (http://www.R-project.org/). *P* values were calculated from one-tailed Student’s *t* test in this study.

## ACKNOWLEDGMENTS

This work was supported by grants 2019YFA0802804 (L.Y.), 2018YFA0801401 (J.C.) and 2018YFC1004602 (J.C.) from MoST, 31925011 (L.Y.), 91940306 (L.Y.), 31822016 (J.C.) and 81872305 (J.C.) from NSFC. We thank Molecular and Cell Biology Core Facility, School of Life Science and Technology, ShanghaiTech University for providing experimental service.

## AUTHOR CONTRIBUTIONS

J. C., L.Y. and X.H. conceived, designed and supervised the project. R.G. and X.L. performed most experiments with the help of G.L., L.W. and J.Wu on cell culture and plasmid construction. J.Wei prepared libraries for DNA sequencing and Z-C.F. and Y.W. performed bioinformatics analyses with the help of W.X., supervised by L.Y. J.C. and L.Y. wrote the paper with inputs from the authors. J.C. managed the project.

## SUPPLEMENTARY FIGURES AND LEGENDS

**Supplementary Fig. 1.**
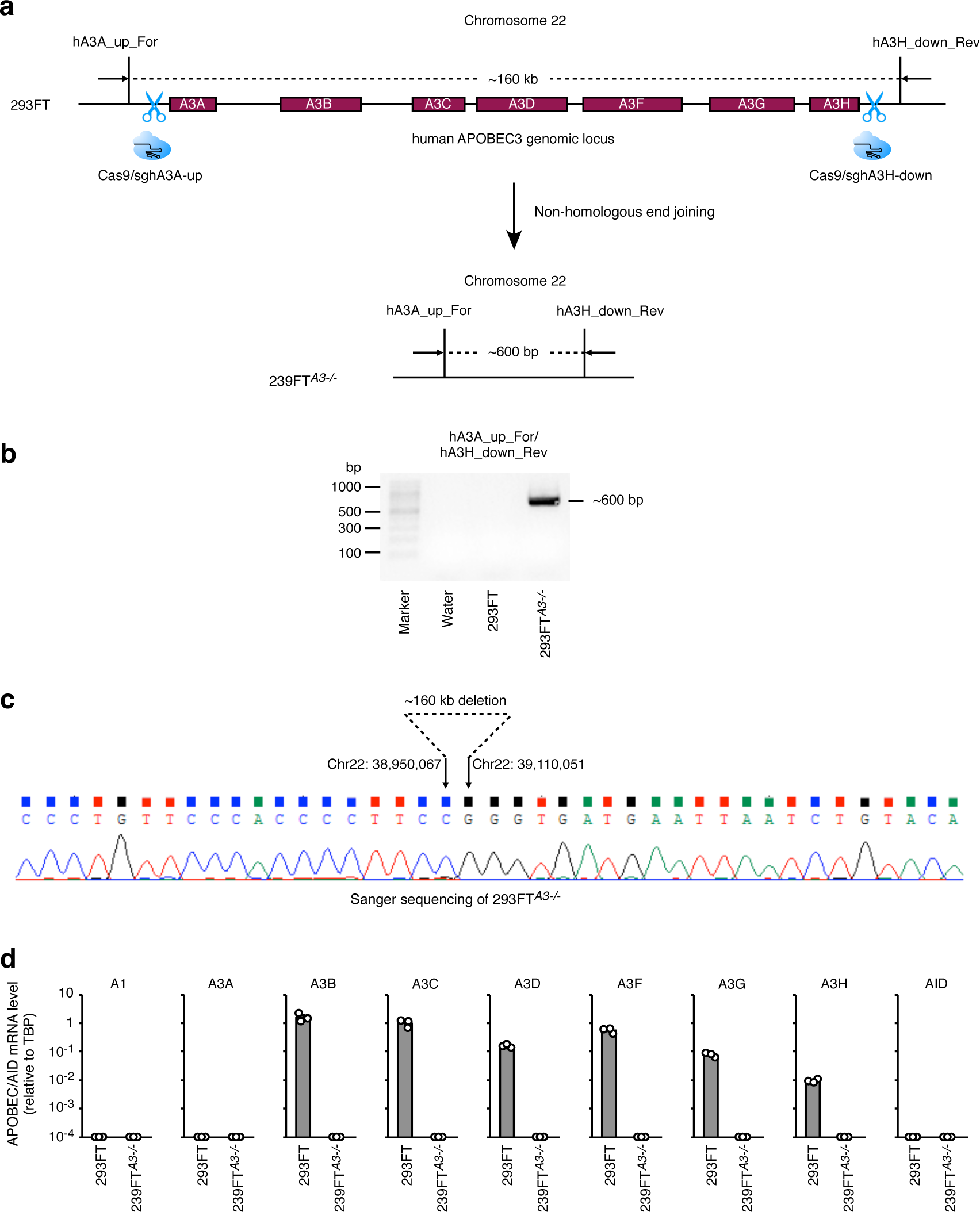
Generation of *APOBEC3*-knockout 293FT cell line. (**a**) Schematic diagrams illustrate the strategy to delete *APOBEC3* gene cluster from the genome of 293FT cells. (**b**) Validation of *APOBEC3*-knockout by RT-PCR. The non-homologous end joining of two Cas9 cleavage site results in a PCR-detectable band (∼600 bp). (**c**) Validation of *APOBEC3*-knockout by Sanger sequencing of the PCR amplified band shown in (**b**). (**d**) Relative APOBEC/AID mRNA expression levels in *APOBEC3*-knockout cells and wildtype cells. Means ± s.d. were from three independent colonies.

**Supplementary Fig. 2.**
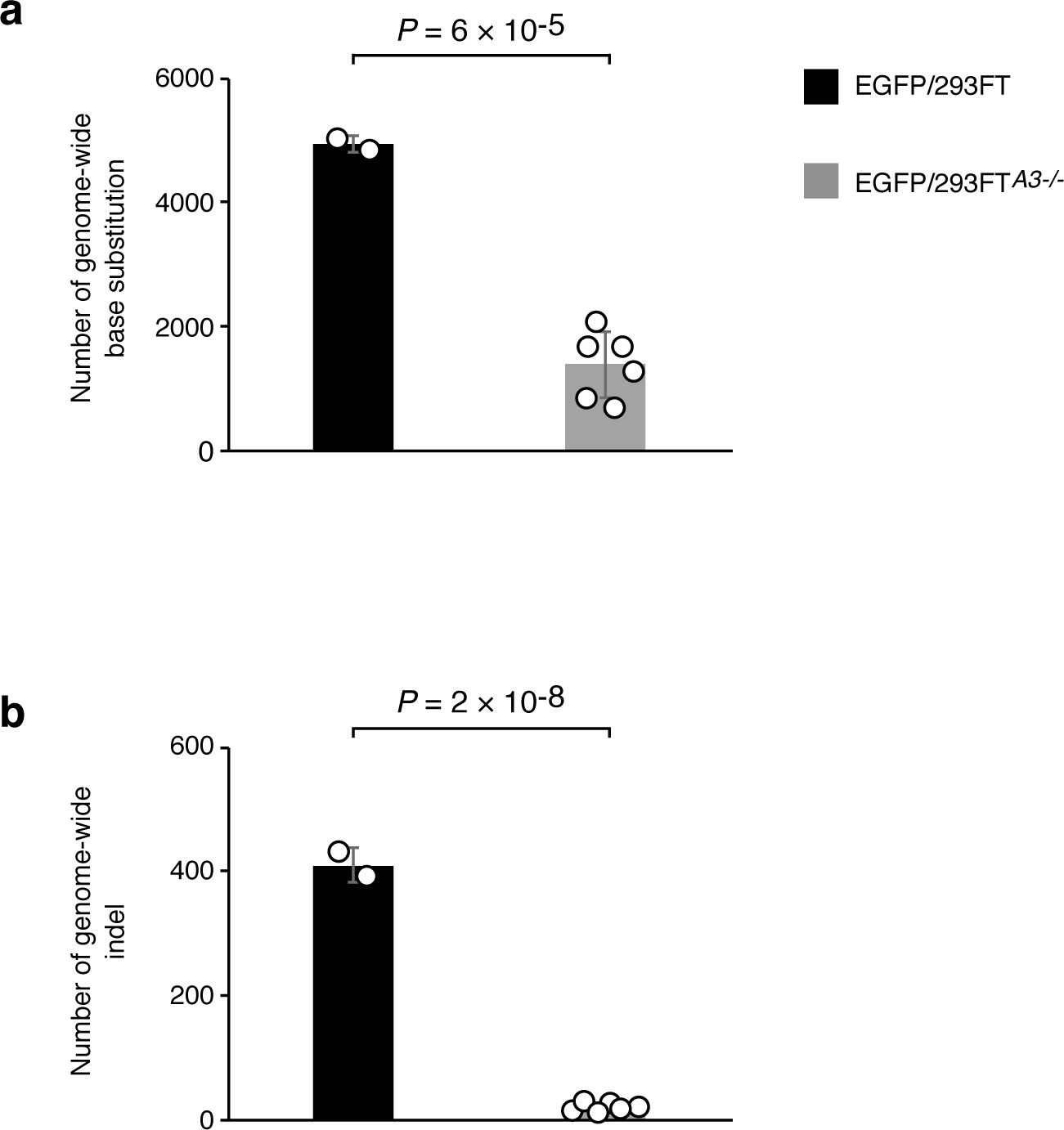
Knockout of *APOBEC3* gene cluster reduced genomic mutation background. (**a**) Comparison of numbers of genome-wide base substitutions in WT 293FT and 293FT*^A3-/-^* cells transfected with EGFP expression plasmid. (**b**) Comparison of numbers of genome-wide indels in WT 293FT and 293FT*^A3-/-^* cells transfected with EGFP expression plasmid. Means ± s.d. were from four independent colonies. *P* value, one-tailed Student’s *t* test.

**Supplementary Fig. 3.**
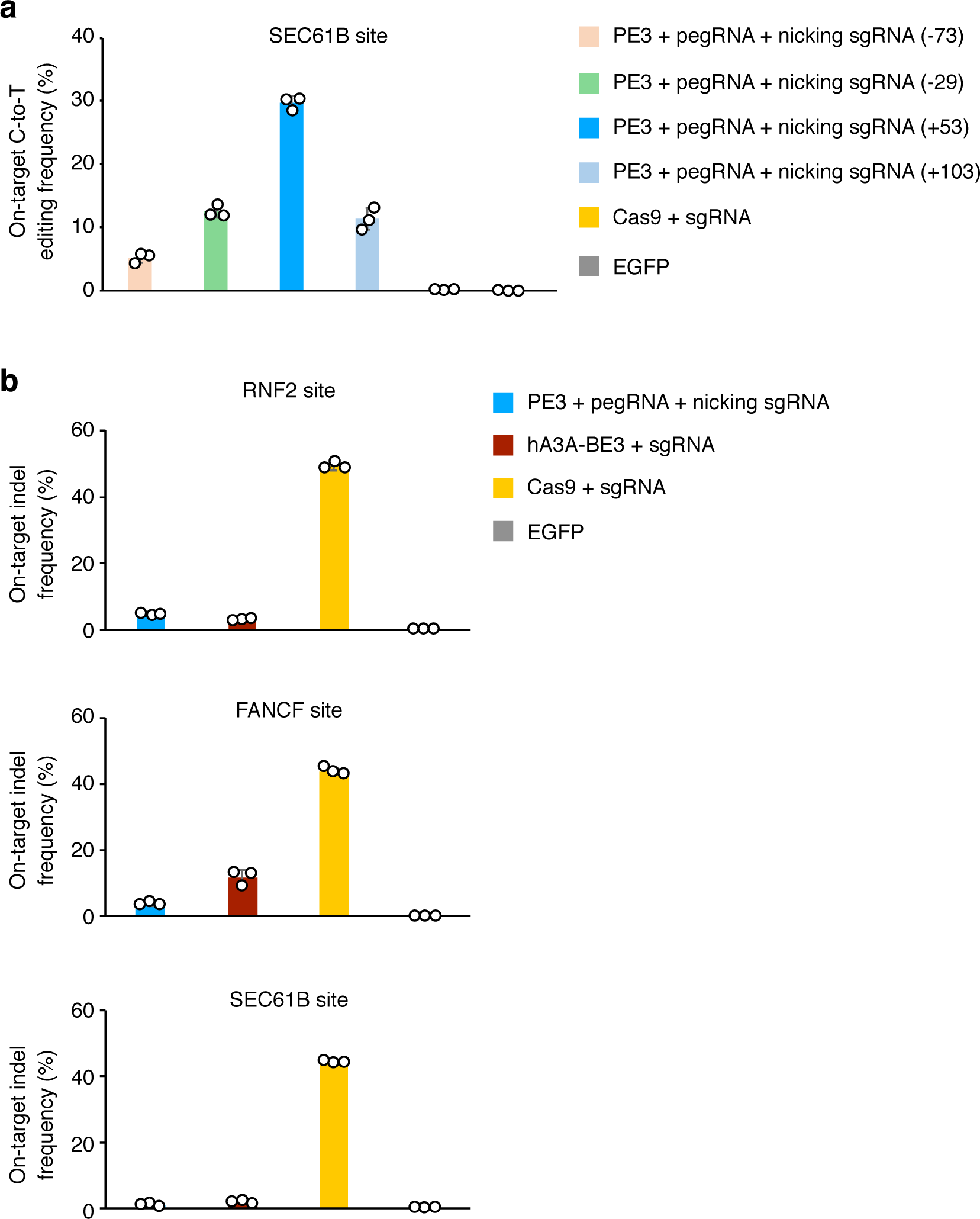
Optimization of nicking gRNA for generating targeted base substitutions at *SEC61B* site and unintended on-target indel frequency. (**a**) On-target C-to-T editing frequencies induced by PE3, pegRNA and different nicking positions, with Cas9 and EGFP as control. (**b**) On-target unintended indel frequencies induced by PE3, hA3A-BE3, Cas9 and EGFP. Means ± s.d. were from three independent colonies.

**Supplementary Fig. 4.**
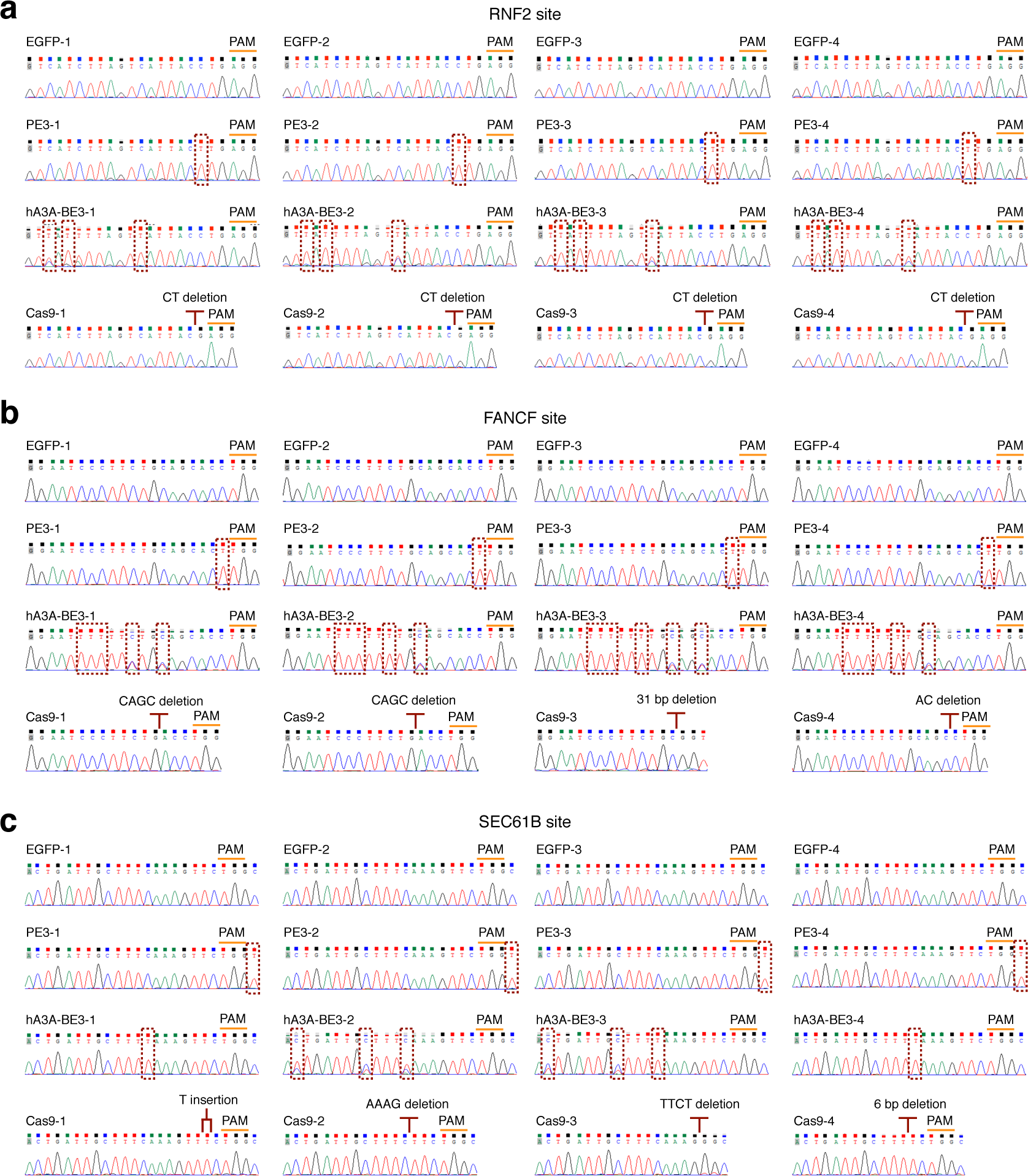
Sanger sequencing of on-target sites in single cell colonies containing intended base substitutions. Sanger sequencing of on-target sites in single cell colonies treated with EGFP, PE3, hA3A-BE3 and Cas9 at *RNF2* (**a**), *FANCF* (**b**) and *SEC61B* (**c**) target sites.

**Supplementary Fig. 5.**
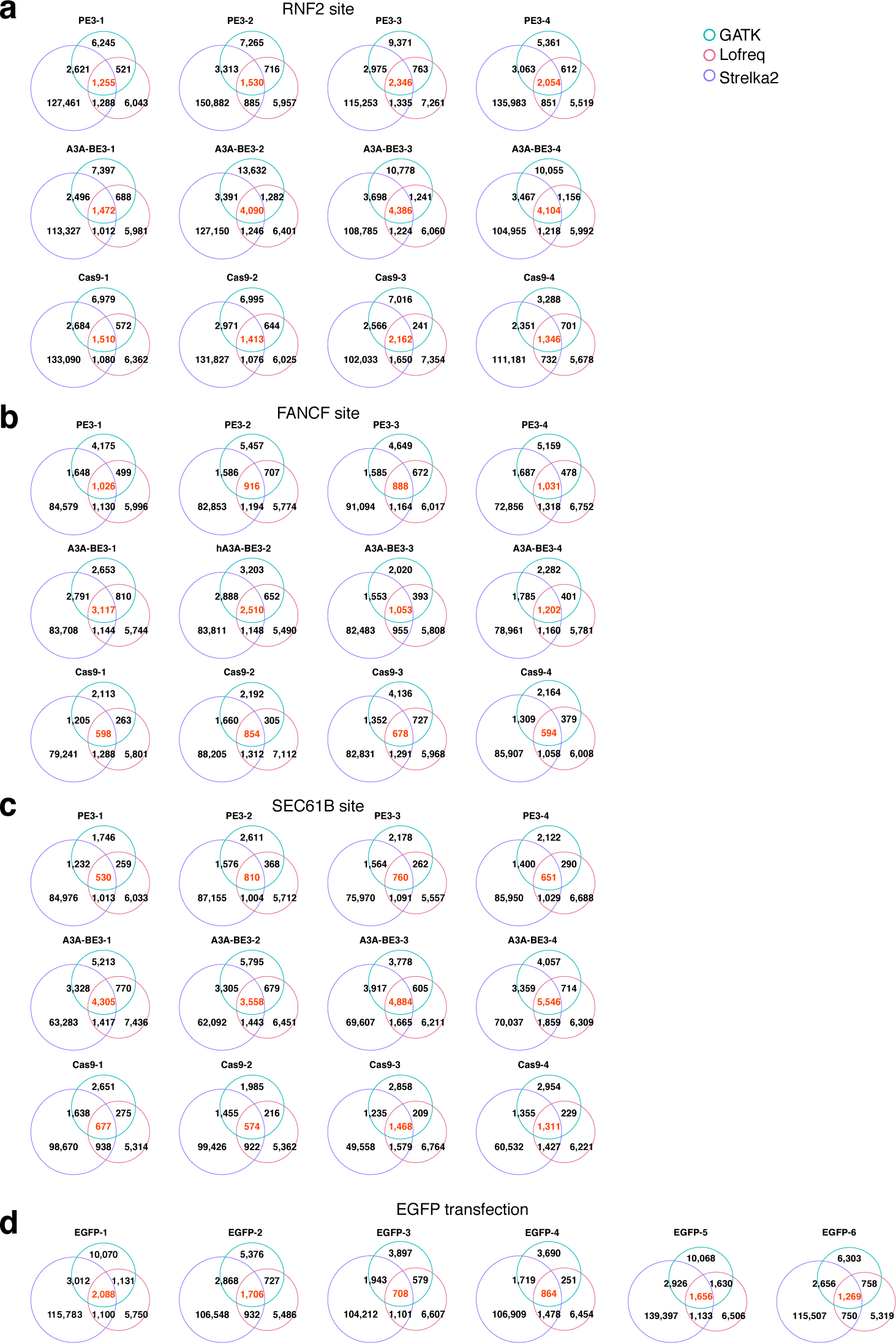
Venn diagrams of genome-wide base substitutions induced by genome editors when generating targeted base substitutions. Venn diagrams of genome-wide base substitutions in single cell colonies detected by three callers after PE3, hA3A-BE3 and Cas9 treatment at *RNF2* (**a**), *FANCF* (**b**) and *SEC61B* (**c**) target sites. (**d**) Venn diagrams of genome-wide base substitutions detected by three callers in EGFP treated single cell colonies.

**Supplementary Fig. 6.**
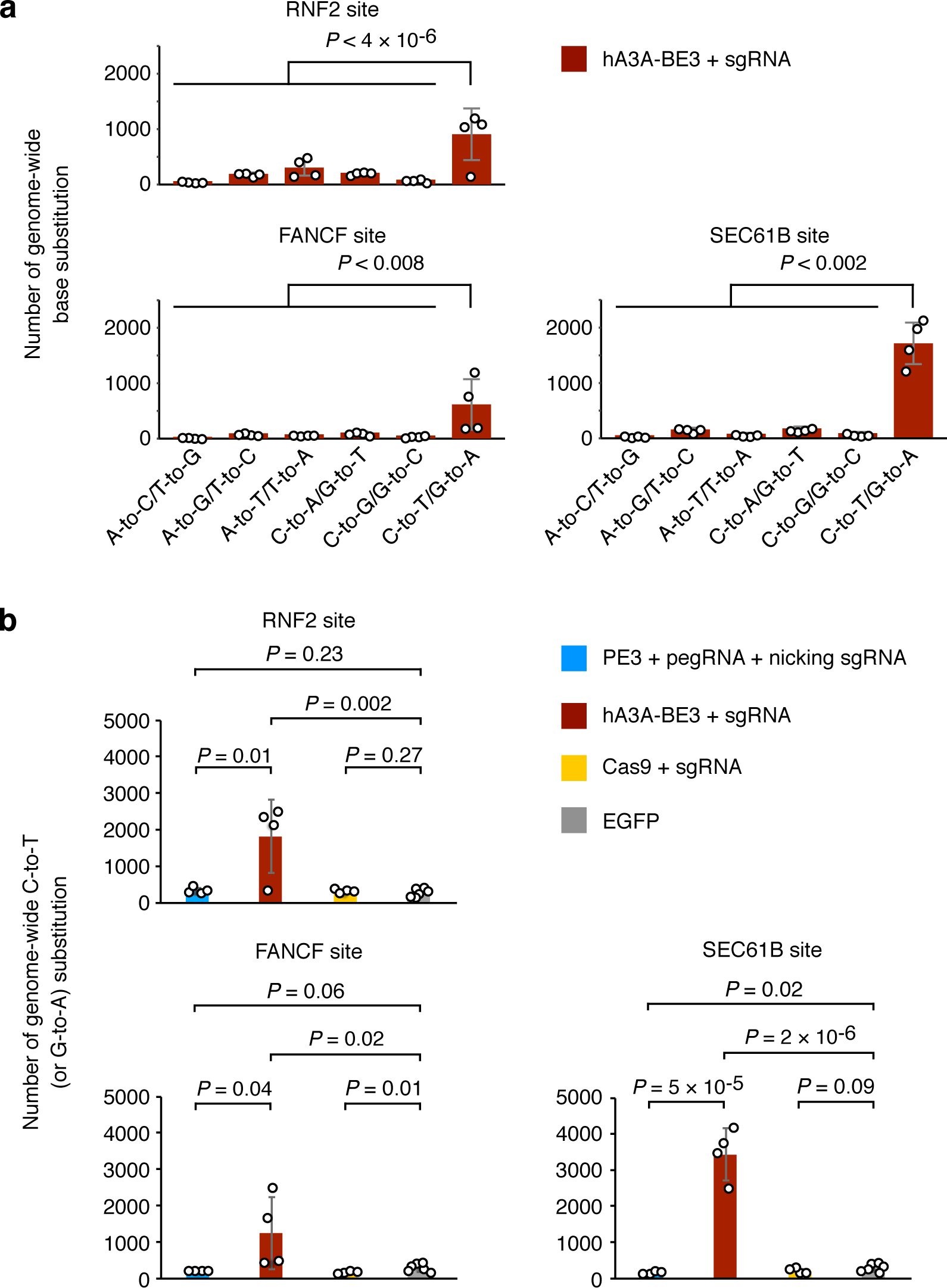
hA3A-BE3 induced genome-wide C-to-T or A-to-G mutations. (**a**) Histograms to show numbers of all six types of genome-wide base substitutions from cells treated with hA3A-BE3. (**b**) Numbers of genome-wide C-to-T/A-to-G substitutions from cells treated with EGFP, Cas9, hA3A-BE3 and PE3. Means ± s.d. were from four independent colonies.

**Supplementary Fig. 7.**
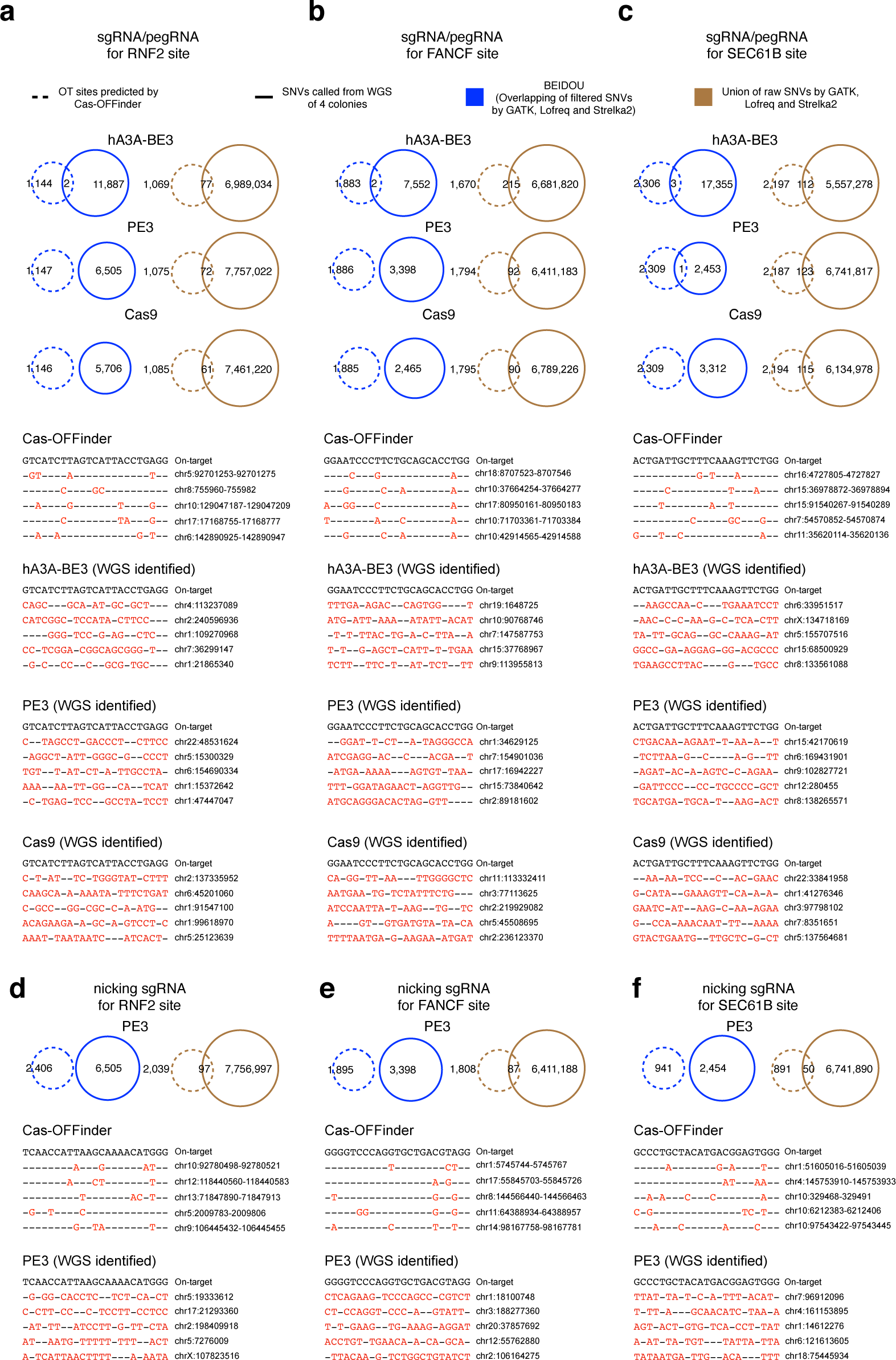
Few overlap of OT sites predicted by Cas-OFFinder and identified by WGS. (**a**-**c**) Comparison of the OT sites predicted by Cas-OFFinder based on pegRNA spacer regions for *RNF2*, *FANCF* and *SEC61B* sites and the OT sites identified by WGS. (**d**-**f**) Comparison of the OT sites predicted by Cas-OFFinder based on the nicking gRNA spacer regions for *RNF2*, *FANCF* and *SEC61B* sites and the OT sites identified by WGS. The on-target sequences were in black and the mismatched bases at OT sites were in red. The intersection (blue circle) or union (brown circle) of callers were used to identify SNVs from WGS data.

**Supplementary Fig. 8.**
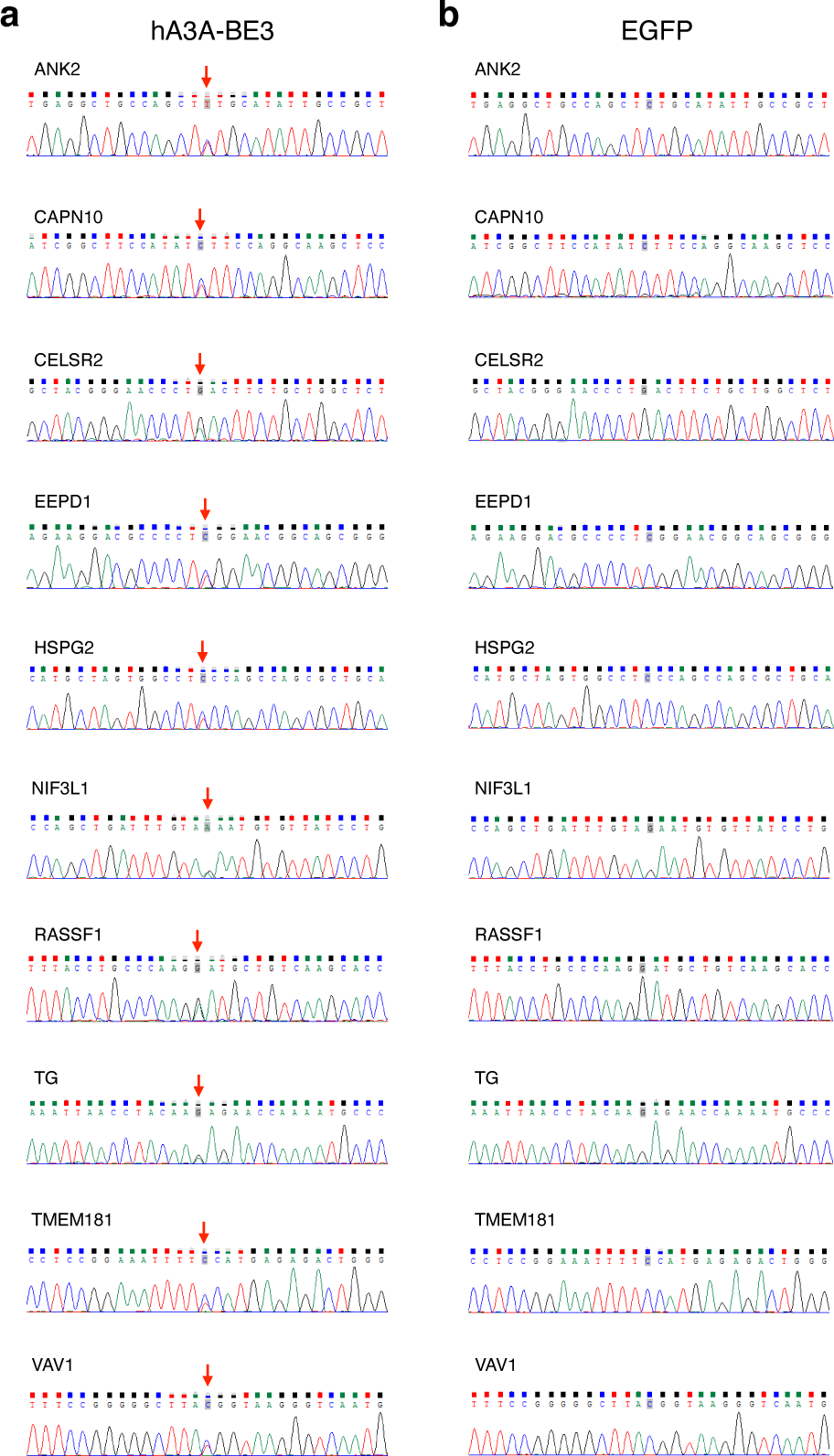
Sanger sequencing of OT site identified by WGS in single cell colonies. Sanger sequencing of OT site identified by WGS in single cell colonies treated with hA3A-BE3 and RNF2-targeting gRNA (**a**) or EGFP (**b**).

**Supplementary Fig. 9.**
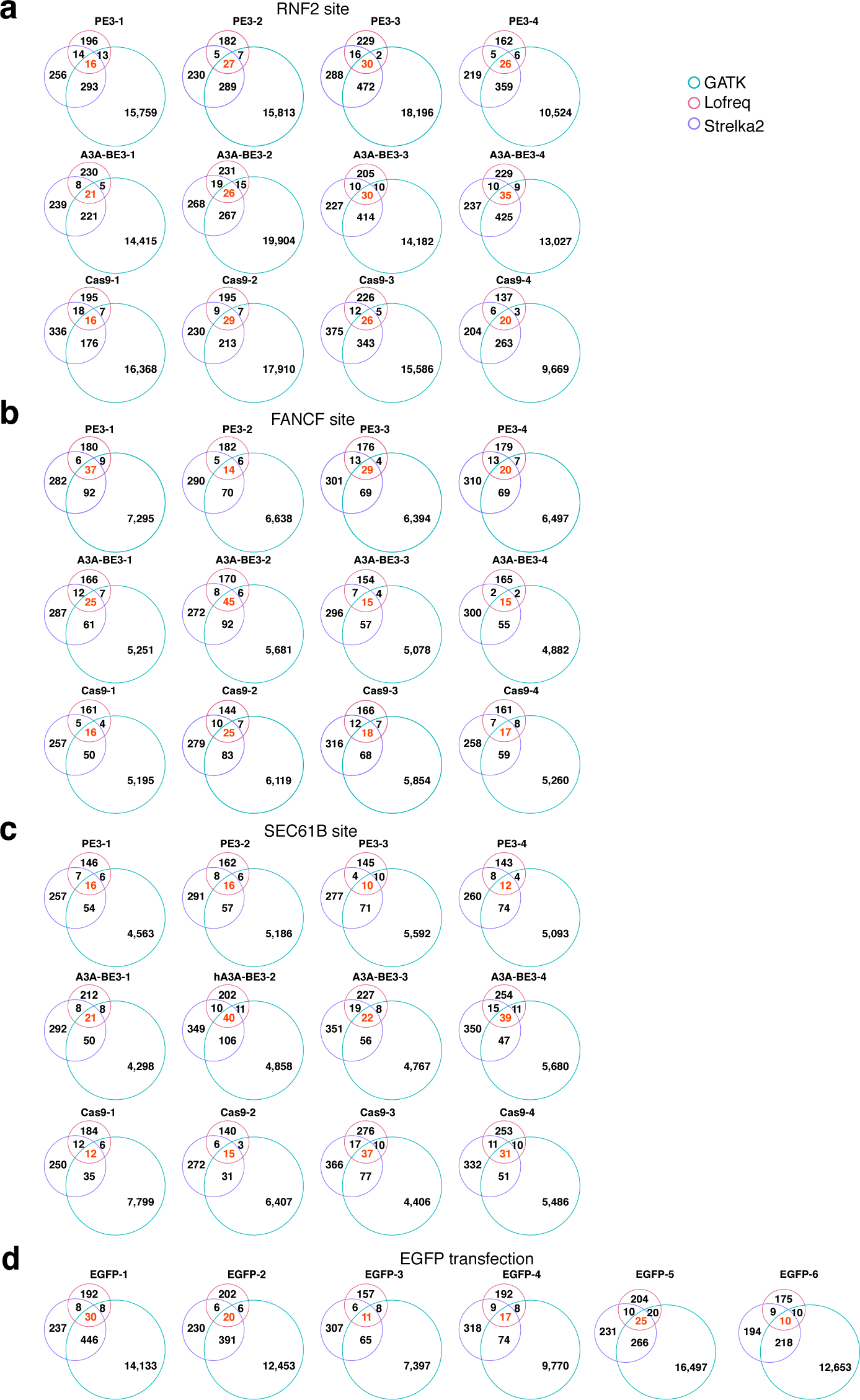
Venn diagrams of genome-wide indels induced by genome editors when generating targeted base substitutions. (**a**) Venn diagrams of genome-wide indels in single cell colonies detected by three callers after PE3, hA3A-BE3 and Cas9 treatment at *RNF2* (**a**), *FANCF* (**b**) and *SEC61B* (**c**) target sites when generating targeted base substitutions. (**d**) Venn diagrams of genome-wide indels detected by three callers in EGFP treated single cell colonies.

**Supplementary Fig. 10.**
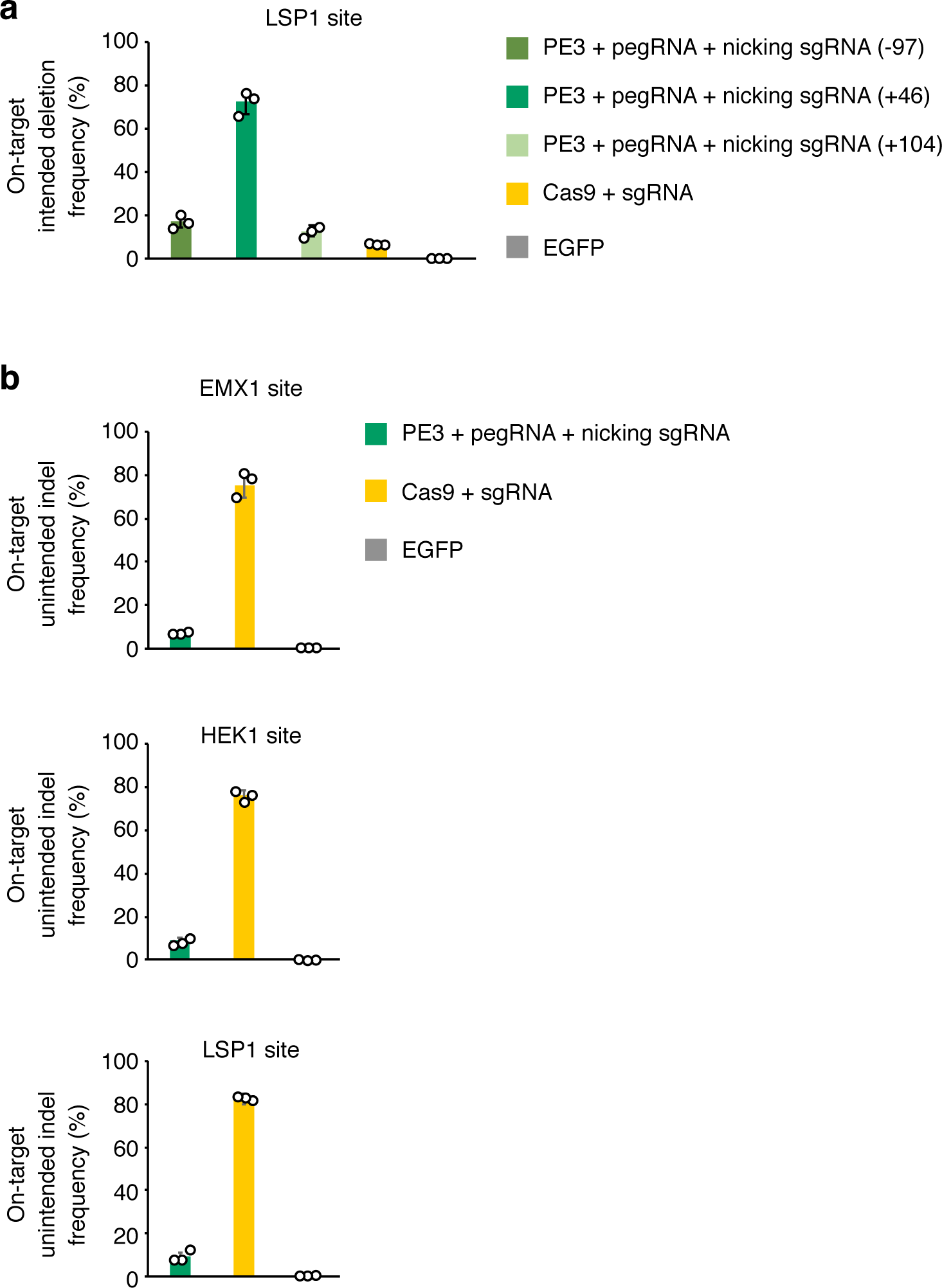
Optimization of nicking gRNA for generating targeted deletions at *LSP1* site and unintended on-target indel frequency. (**a**) On-target intended deletion frequencies induced by PE3, pegRNA and different nicking positions, with Cas9 and EGFP as control. (**b**) On-target unintended indel frequencies by PE3, Cas9 and EGFP. Means ± s.d. were from four independent colonies.

**Supplementary Fig. 11.**
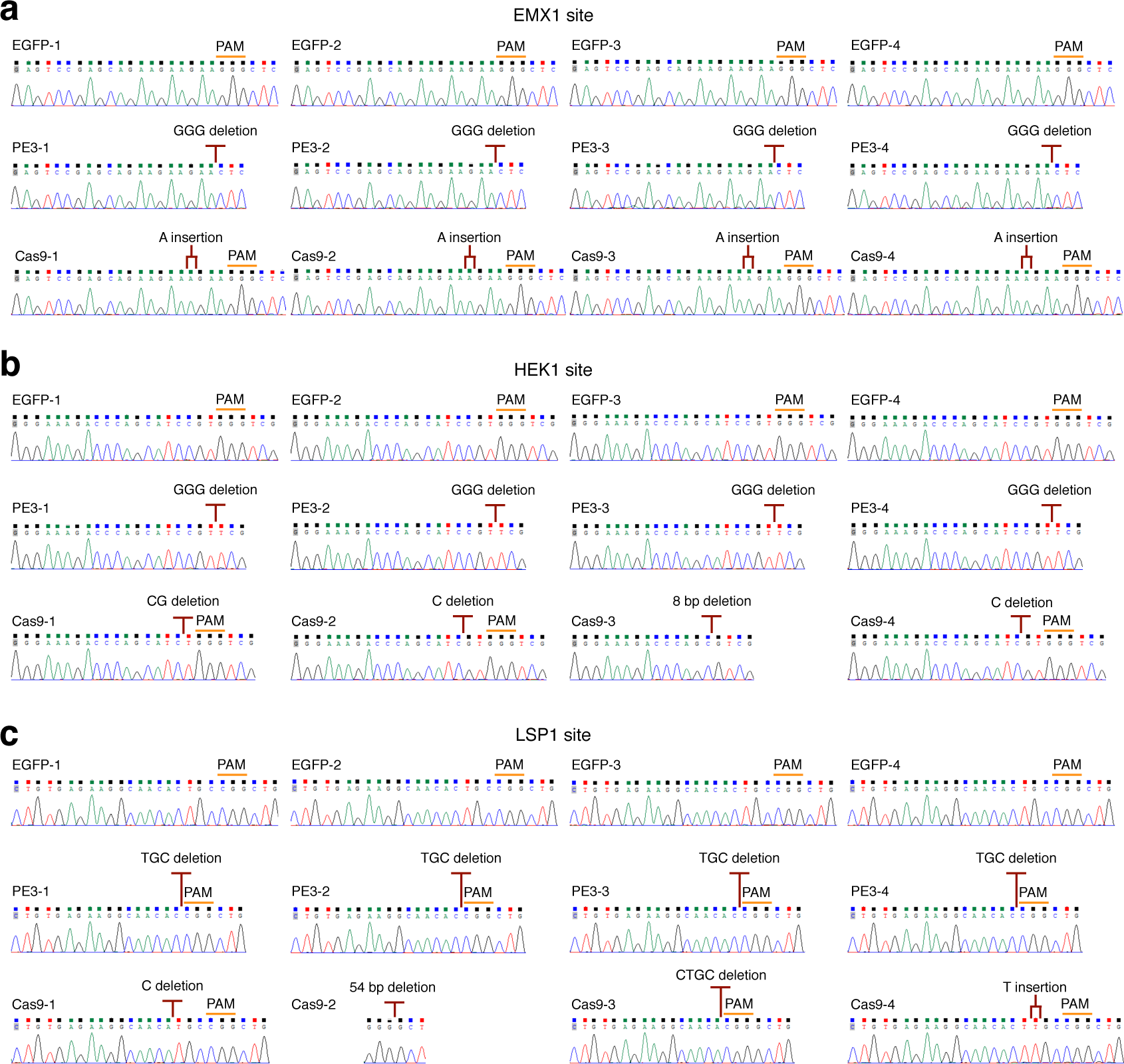
Sanger sequencing of on-target sites in single cell colonies containing intended deletions. Sanger sequencing of on-target sites in single cell colonies after EGFP, PE3, Cas9 treatment at *EMX1* (**a**), *HEK1* (**b**), *LSP1* (**c**) target sites when generating intended deletions.

**Supplementary Fig. 12.**
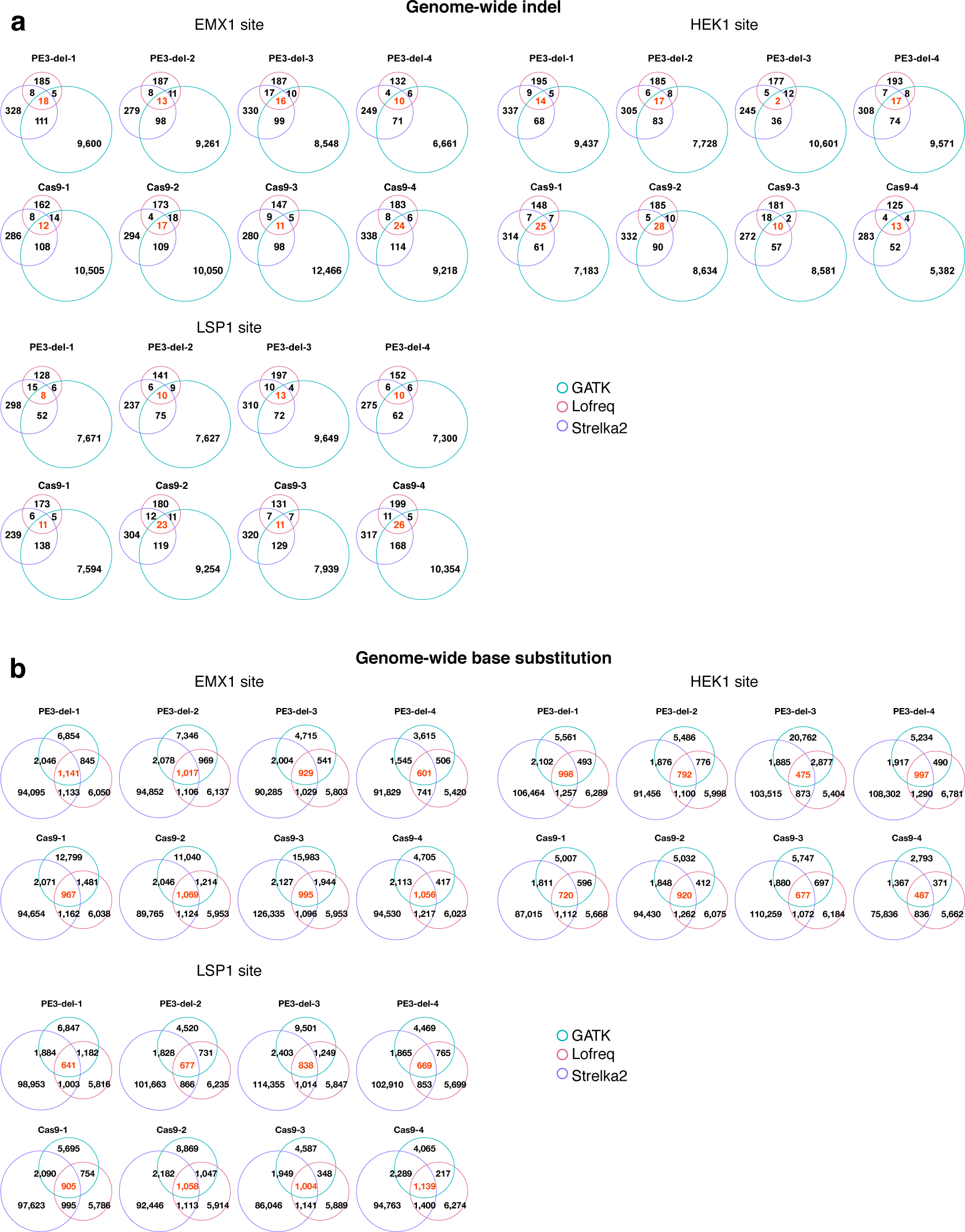
Venn diagrams of genome-wide indels and base substitutions induced by genome editors when generating targeted deletions. (**a**) Venn diagrams of genome-wide indels in single cell colonies detected by three callers after PE3 and Cas9 treatment at *EMX1, HEK1*, *LSP1* target sites when generating targeted deletions. (**b**) Venn diagrams of genome-wide base substitutions in single cell colonies detected by three callers after PE3 and Cas9 treatment at *EMX1, HEK1*, *LSP1* target sites when generating targeted deletions.

**Supplementary Fig. 13.**
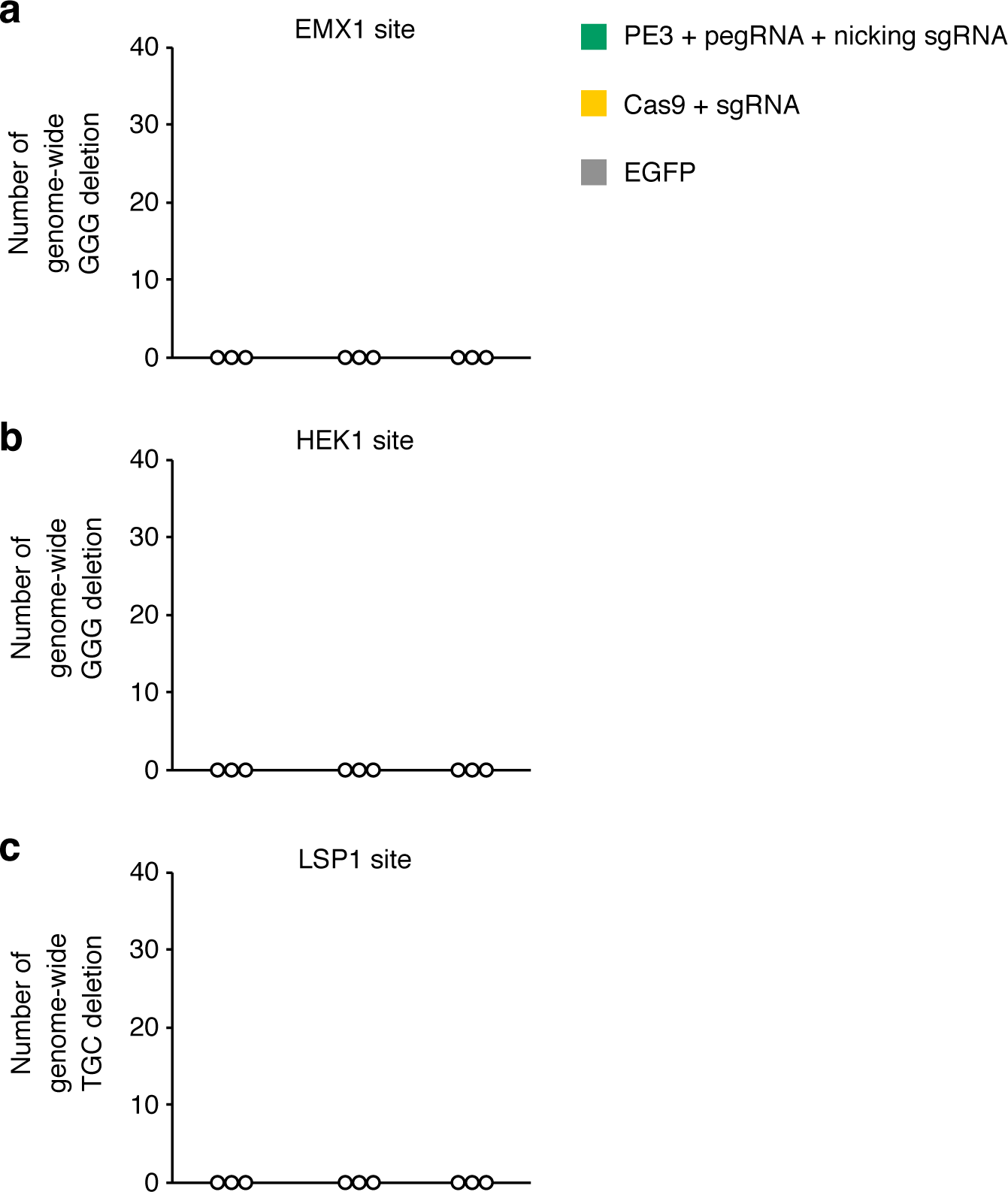
PE3 induced no intended 3-bp deletion on a genome-wide scale. (**a**) Numbers of genome-wide GGG deletions after cells treated with PE3, Cas9 and EGFP at *EMX1* site when generating targeted GGG-deletion. (**b**) Numbers of genome-wide GGG deletions after cells with PE3, Cas9 and EGFP at *HEK1* site when generating targeted GGG-deletion. (**c**) Numbers of genome-wide TGC deletions after cells treated with PE3, Cas9 and EGFP at *LSP1* site when generating targeted TGC deletions.

**Supplementary Fig. 14.**
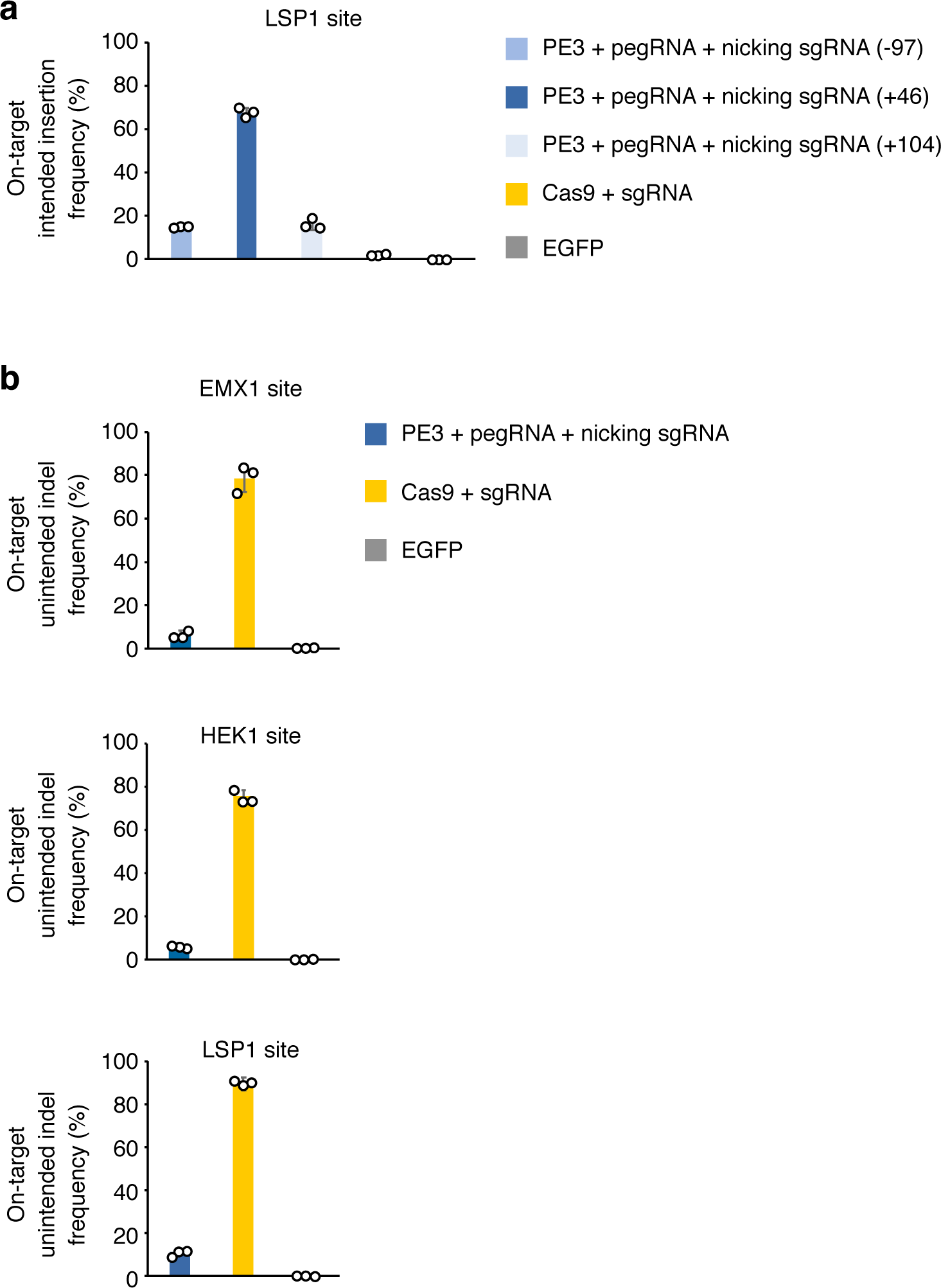
Optimization of nicking gRNA for generating targeted insertions at LSP1 site and unintended on-target indel frequency. (**a**) On-target intended insertion frequencies induced by PE3, pegRNA and different nicking positions, with Cas9 and EGFP as control. (**b**) On-target unintended indel frequencies by PE3, Cas9 and EGFP at three target sites. Means ± s.d. were from four independent colonies.

**Supplementary Fig. 15.**
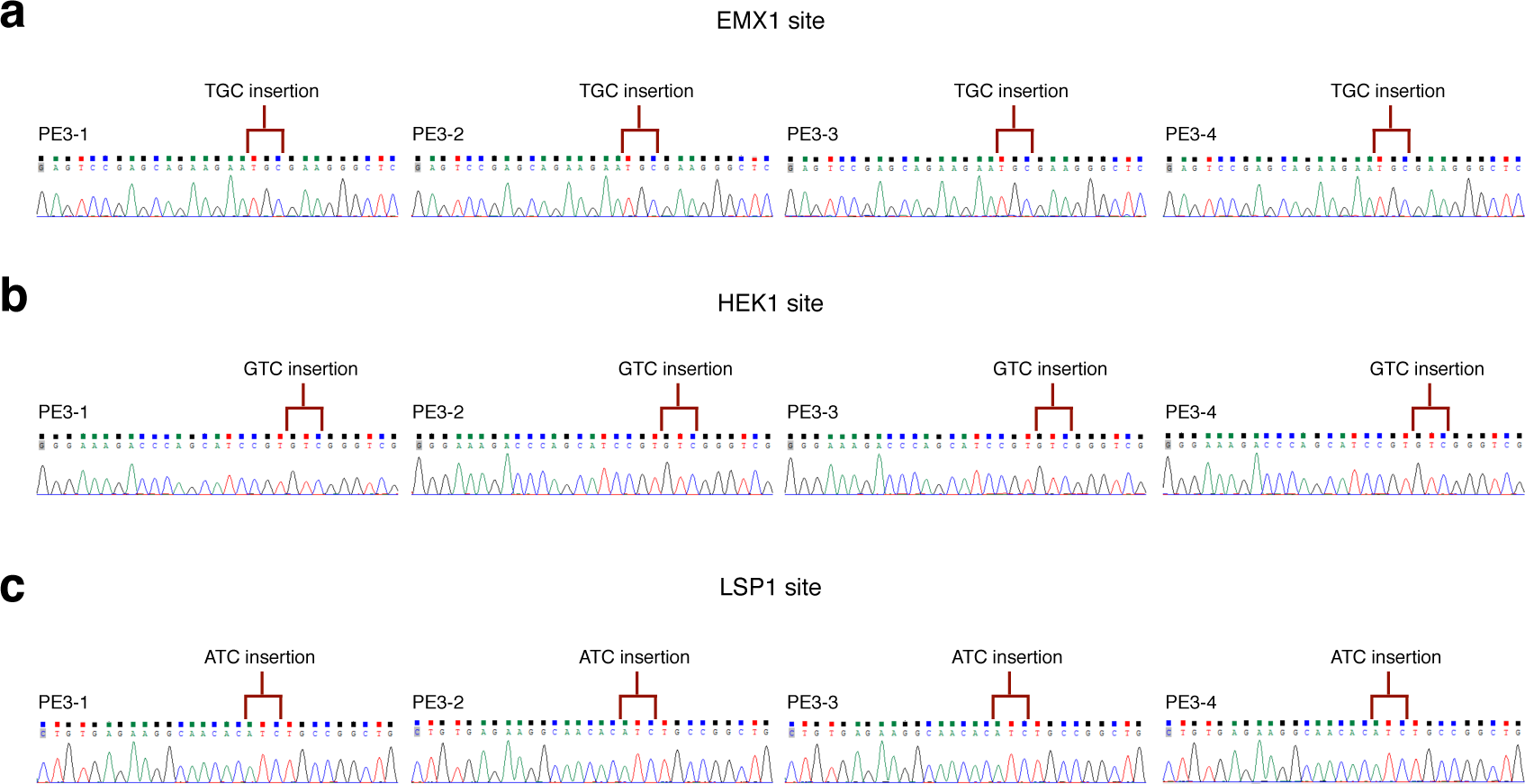
Sanger sequencing of on-target sites in single cell colonies containing intended insertions. (**a**) Sanger sequencing validation of on-target site in single cell colonies containing intended TGC insertions after PE3 treatment at *EMX1* site. (**b**) Sanger sequencing validation of on-target site in single cell colonies containing intended GTC insertions after PE3 treatment at *HEK1* site. (**c**) Sanger sequencing validation of on-target site in single cell colonies containing intended ATC insertions after PE3 treatment at *LSP1* site.

**Supplementary Fig. 16.**
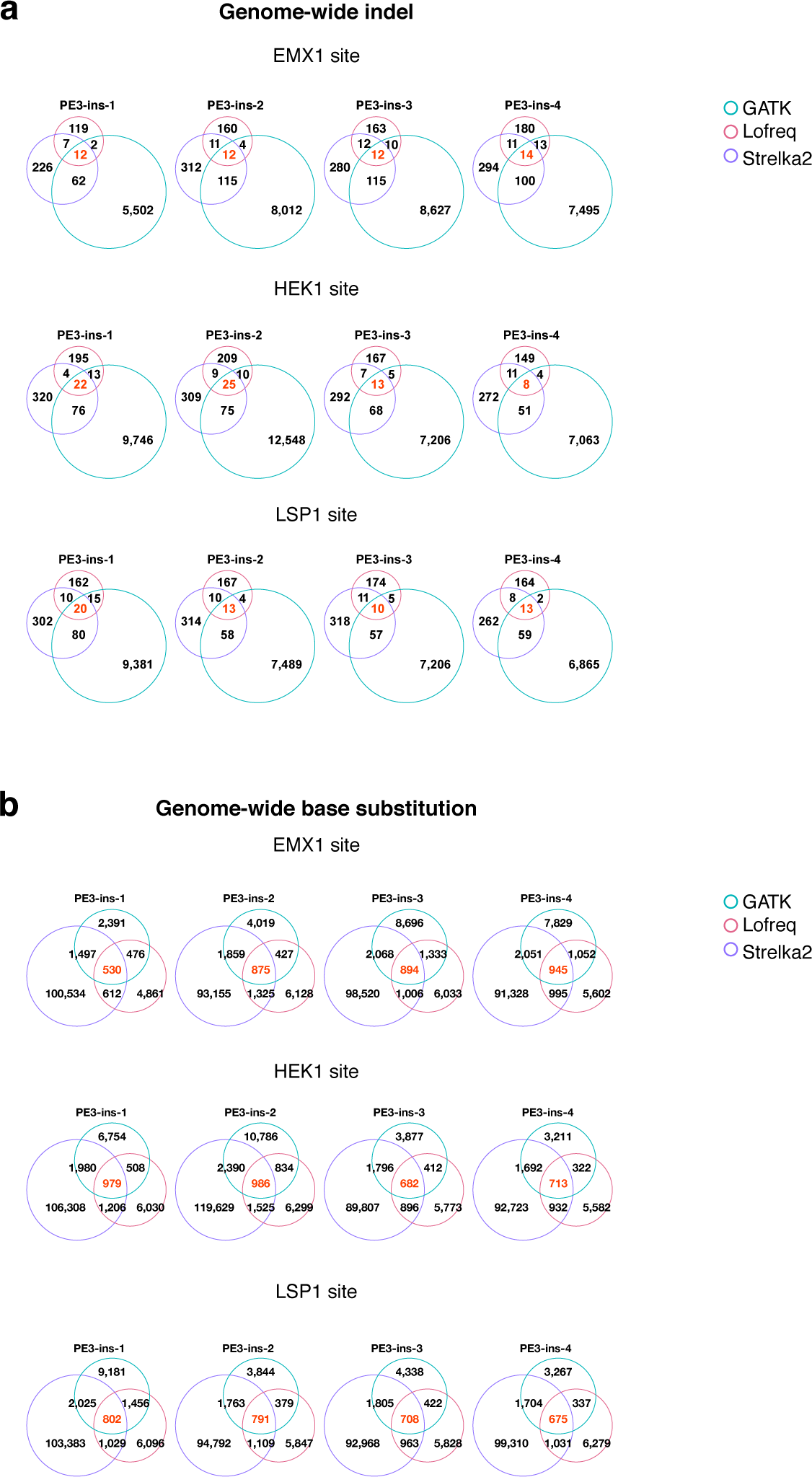
Venn diagrams of genome-wide indels and base substitutions induced by genome editors when generating targeted insertions. (**a**) Venn diagrams of genome-wide indels by three callers in single cell colonies detected after PE3 treatment at *EMX1*, *HEK1*, *LSP1* sites when generating targeted insertions. (**b**) Venn diagrams of genome-wide base substitutions in single cell colonies detected by three callers after PE3 treatment at *EMX1*, *HEK1*, *LSP1* sites when generating targeted insertions.

**Supplementary Fig. 17.**
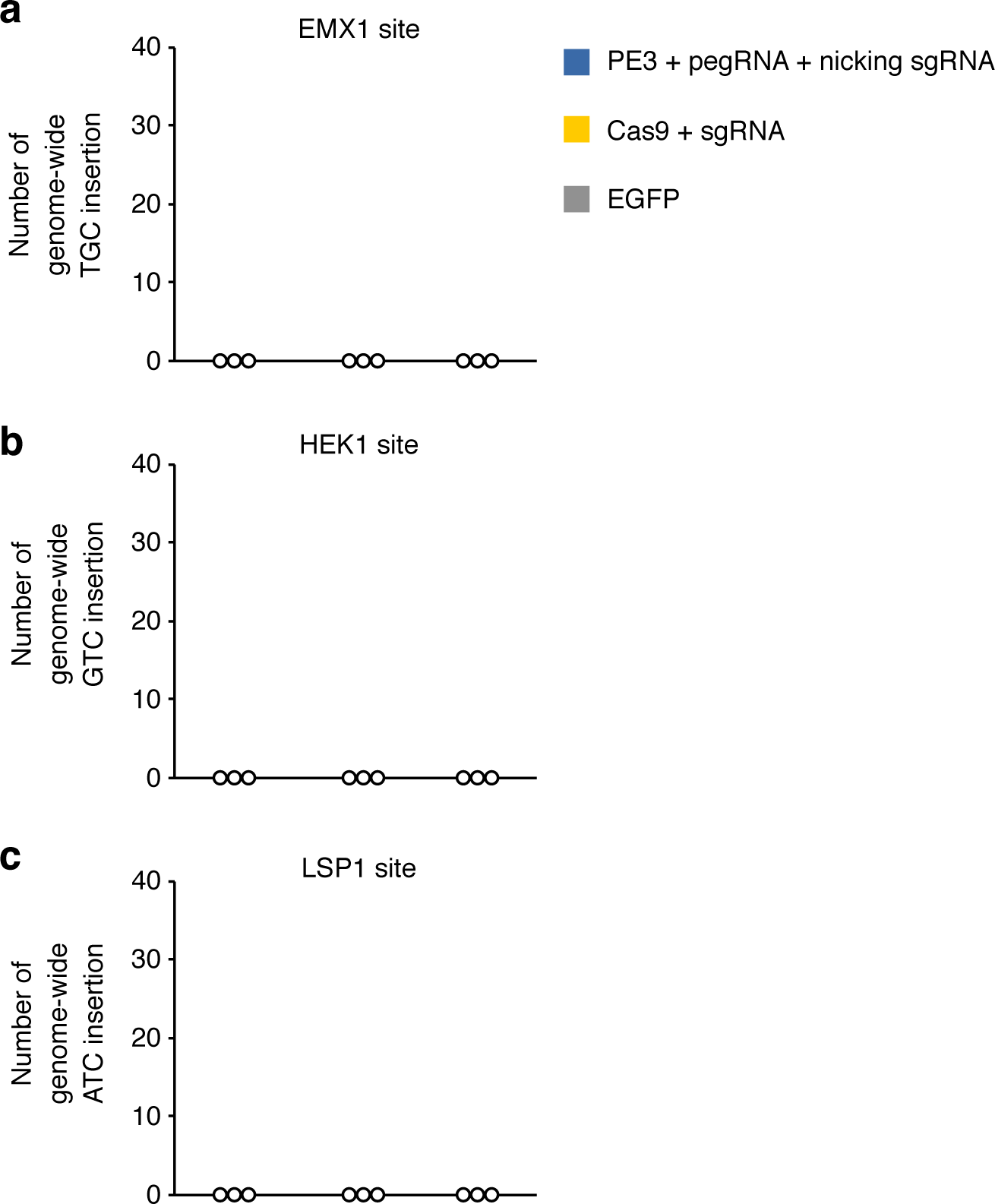
PE3 induced no intended 3-bp insertion on a genome-wide scale. (**a**) Numbers of genome-wide TGC insertions after cells treated with PE3, Cas9 and EGFP at *EMX1* site. (**b**) Numbers of genome-wide GTC insertions after cells treated with PE3, Cas9 and EGFP at *HEK1* site. (**c**) Numbers of genome-wide ATC insertions after cells treated with PE3, Cas9 and EGFP at *LSP1* site.

**Supplementary Fig. 18.**
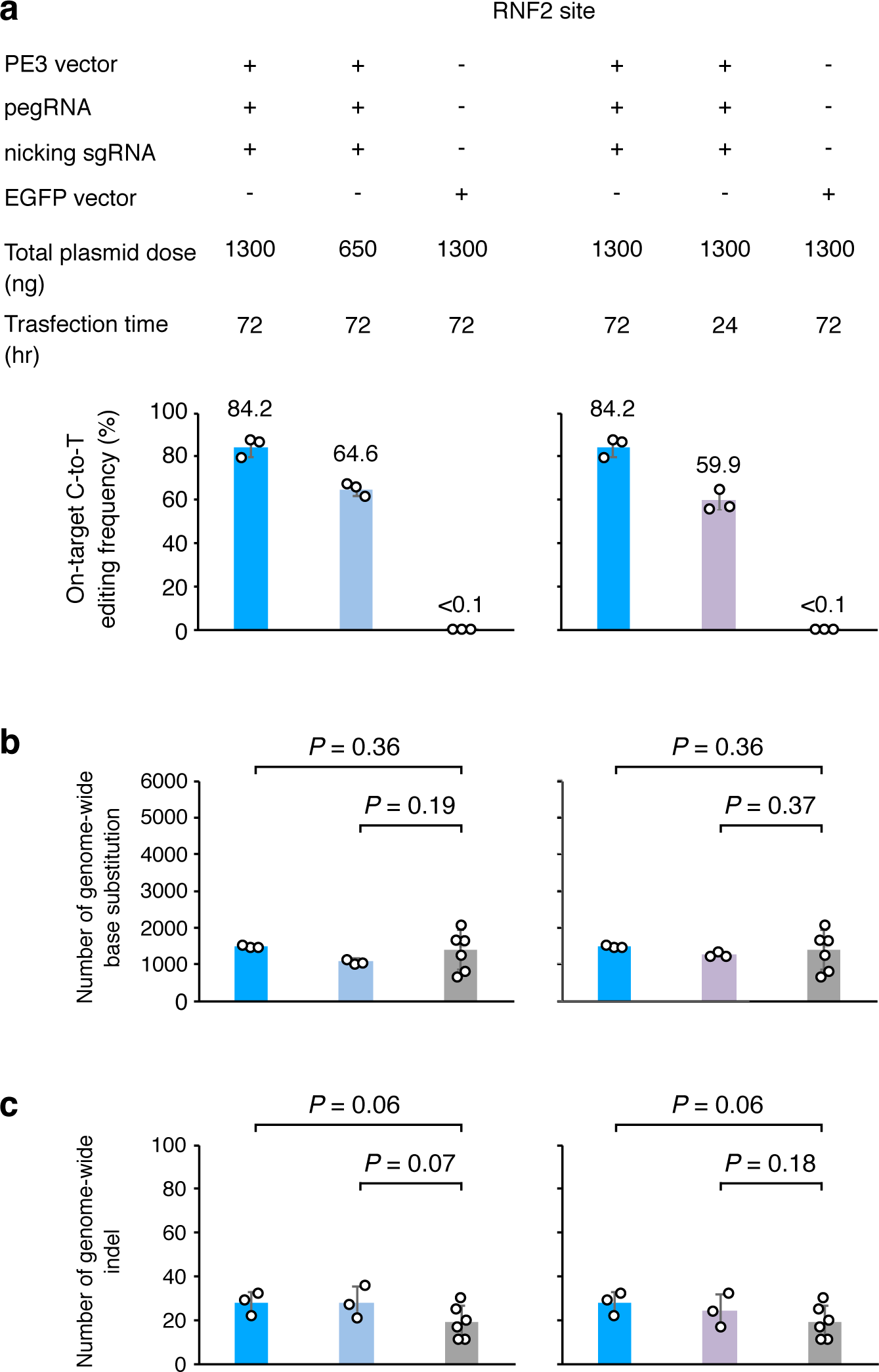
PE3 induced background levels of genome-wide OT mutations via plasmid delivery of different doses and time periods. (**a**) On-target C-to-T editing frequencies induced by PE3 via plasmid delivery of different doses and time periods. (**b**) Numbers of genome-wide base substitutions induced by PE3 via plasmid delivery of different doses and time periods. (**c**) Numbers of genome-wide indels induced by PE3 via plasmid delivery of different doses and time periods. The data for Cas9 and EGFP in (**a**-**c**) are the same.

**Supplementary Fig. 19.**
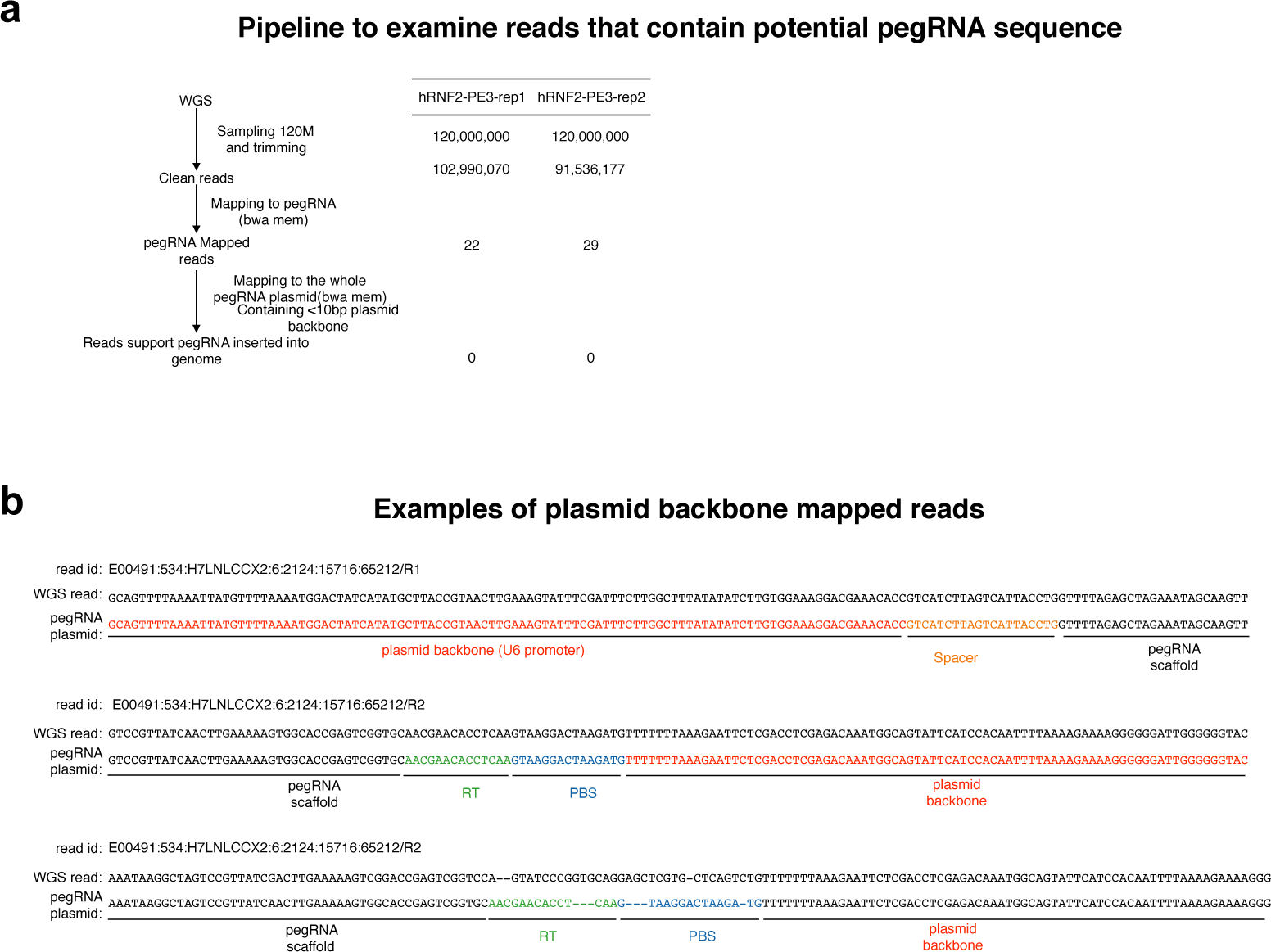
pegRNA backbone sequence was not detected in genomewide OT insertions. (**a**) Examining reads that contains potential pegRNA sequence inserted into genome by aligning raw WGS reads of hRNF3-PE3-rep1 and hRNF3-PE3-rep2 to pegRNA sequence. Left: Schematic overview of the pipeline. Right: count of reads pass each step. (**b**) Examples of pegRNA-containing reads that can be aligned to pegRNA plasmid (vector pU6-pegRNF2) backbone in (**a**), which are not considered as authentic insertions into genome.

**Supplementary Fig. 20.**
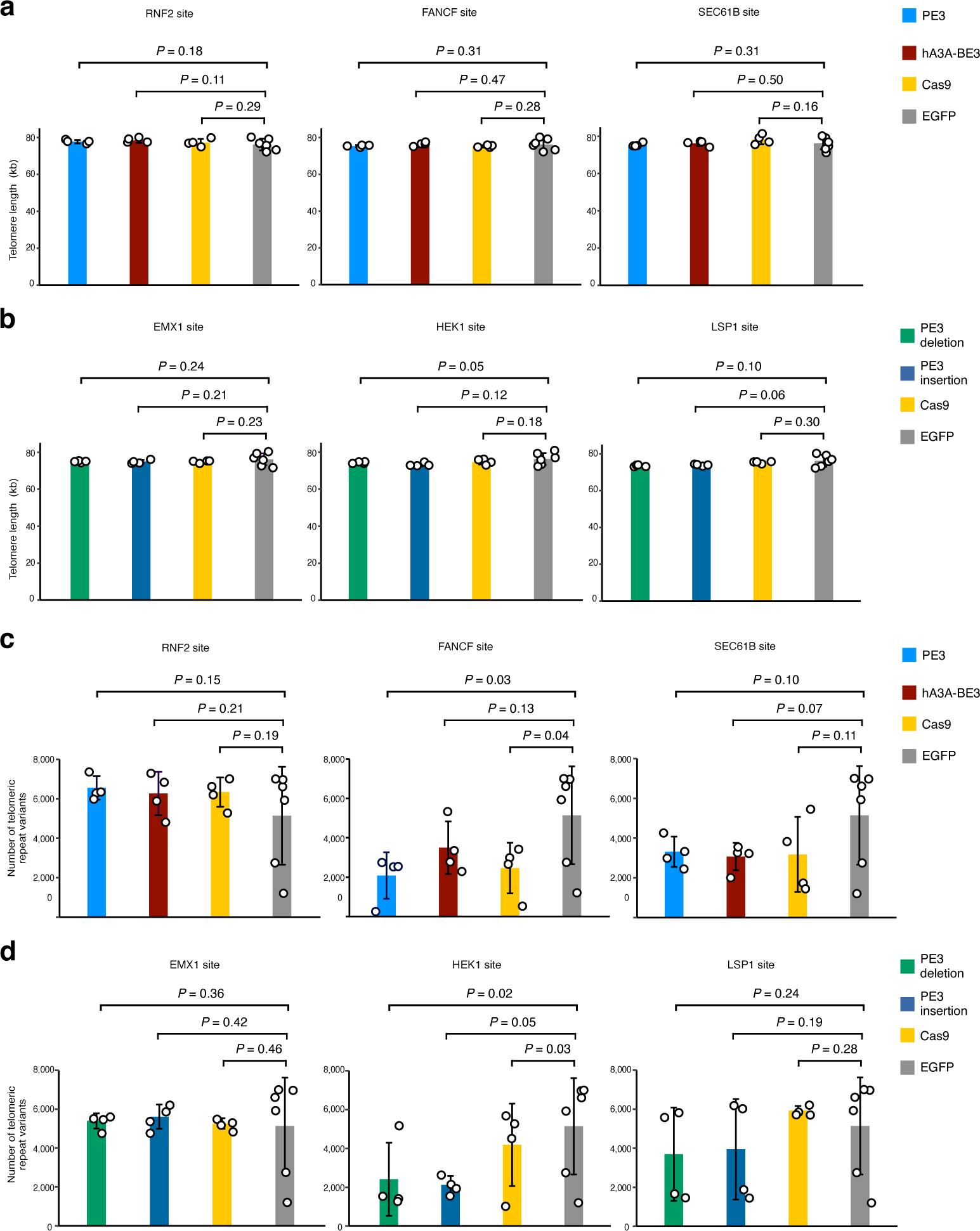
Telomere integrity was not affected by PE3. (**a**, **b**) Telomere lengths in single cell colonies treated by PE3, hA3A-BE3, Cas9 when generating on-target base substitutions (**a**) or indels (**b**) were calculated from the WGS data. (**c**, **d**) The number of telomeric repeat variants in single cell colonies treated by PE3 or Cas9 when generating on-target base substitutions (**c**) or indels (**d**) were calculated from the WGS data. Means ± s.d. were from four independent colonies.

**Supplementary Fig. 21.**
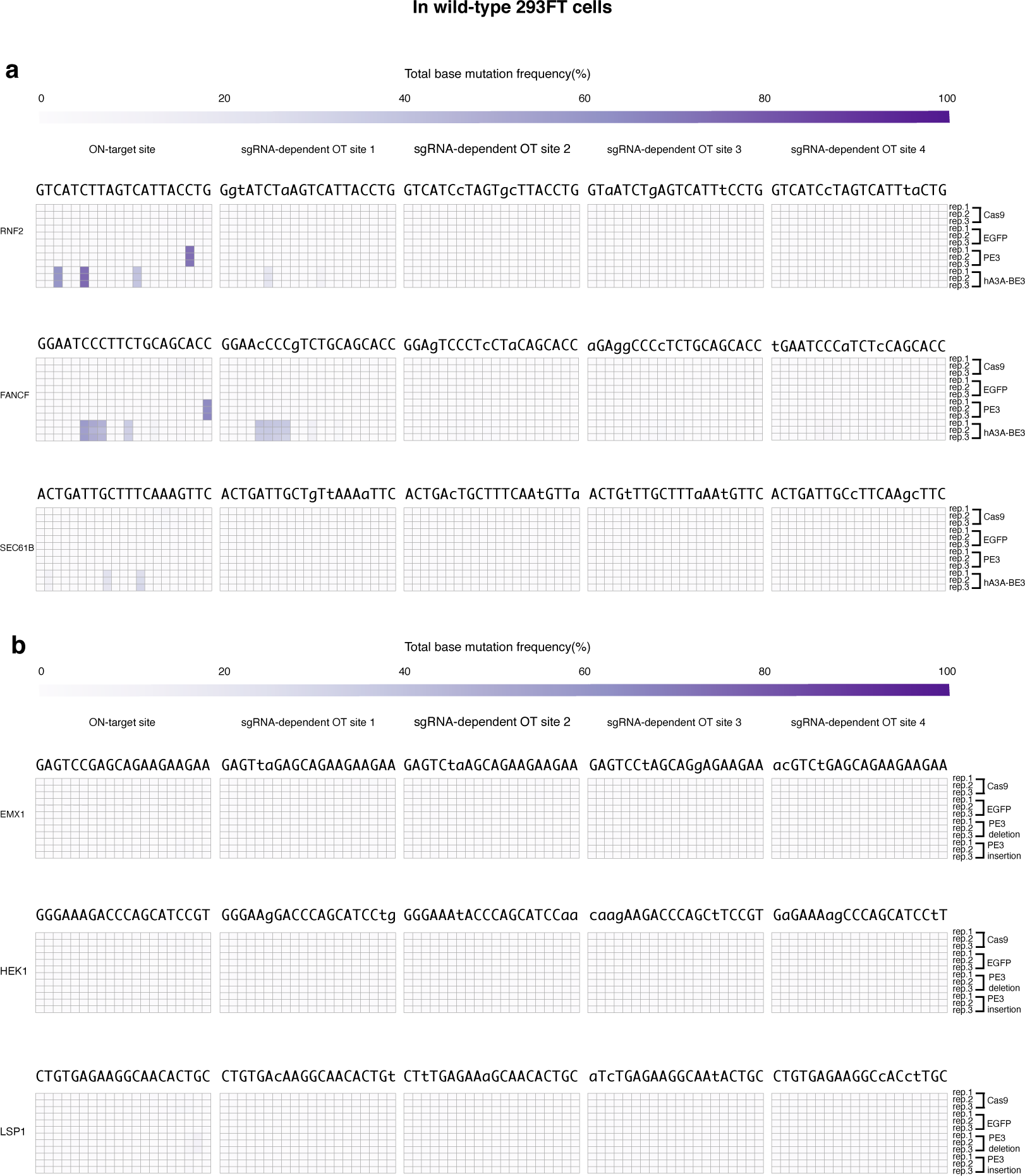
No detectable base substitution induced by PE3 at gRNA-dependent OT sites in wild-type 293FT cells. (**a**) The heatmaps of base substitution frequencies at on-target sites and previously identified or predicted gRNA-dependent OT sites for *RNF2, FANCF, SEC61B* in WT 293FT cells. (**b**) The heatmaps of base substitution frequencies at on-target sites and previously identified or predicted gRNA-dependent OT sites for *EMX1, HEK1, LSP* in WT 293FT cells.

**Supplementary Fig. 22.**
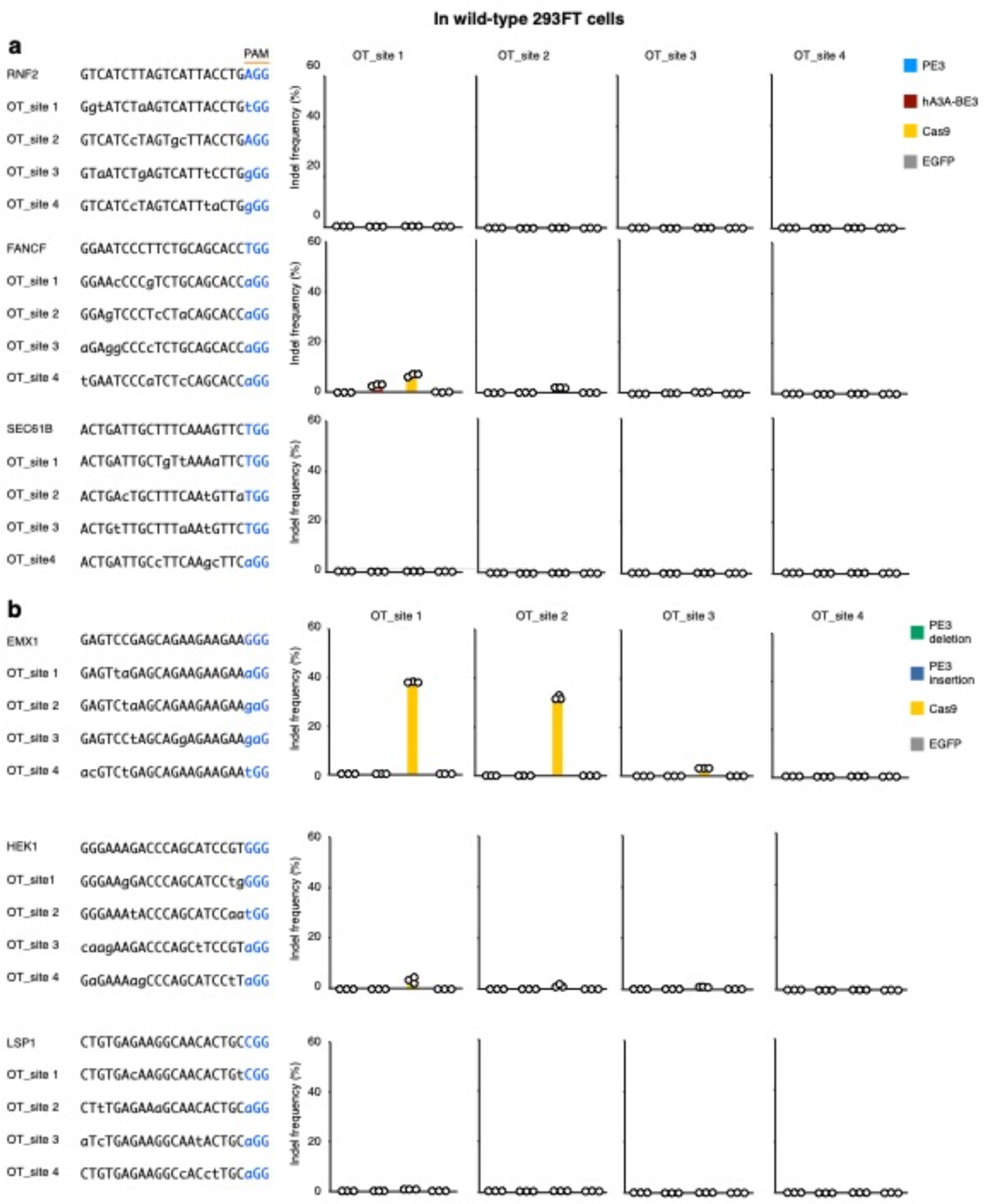
No detectable indel induced by PE3 at gRNA-dependent OT sites in wild-type 293FT cells. (**a**) The sequences of on-target and previously identified or predicted gRNA-dependent OT sites for *RNF2, FANCF, SEC61B* (left) and the indel frequencies at those OT sites (right). (**b**) The sequences of on-target and previously identified or predicted gRNA-dependent OT sites for *EMX1, HEK1, LSP* (left) and the indel frequencies at those OT sites were shown (right). Means ± s.d. were from three independent experiments.

**Supplementary Fig. 23.**
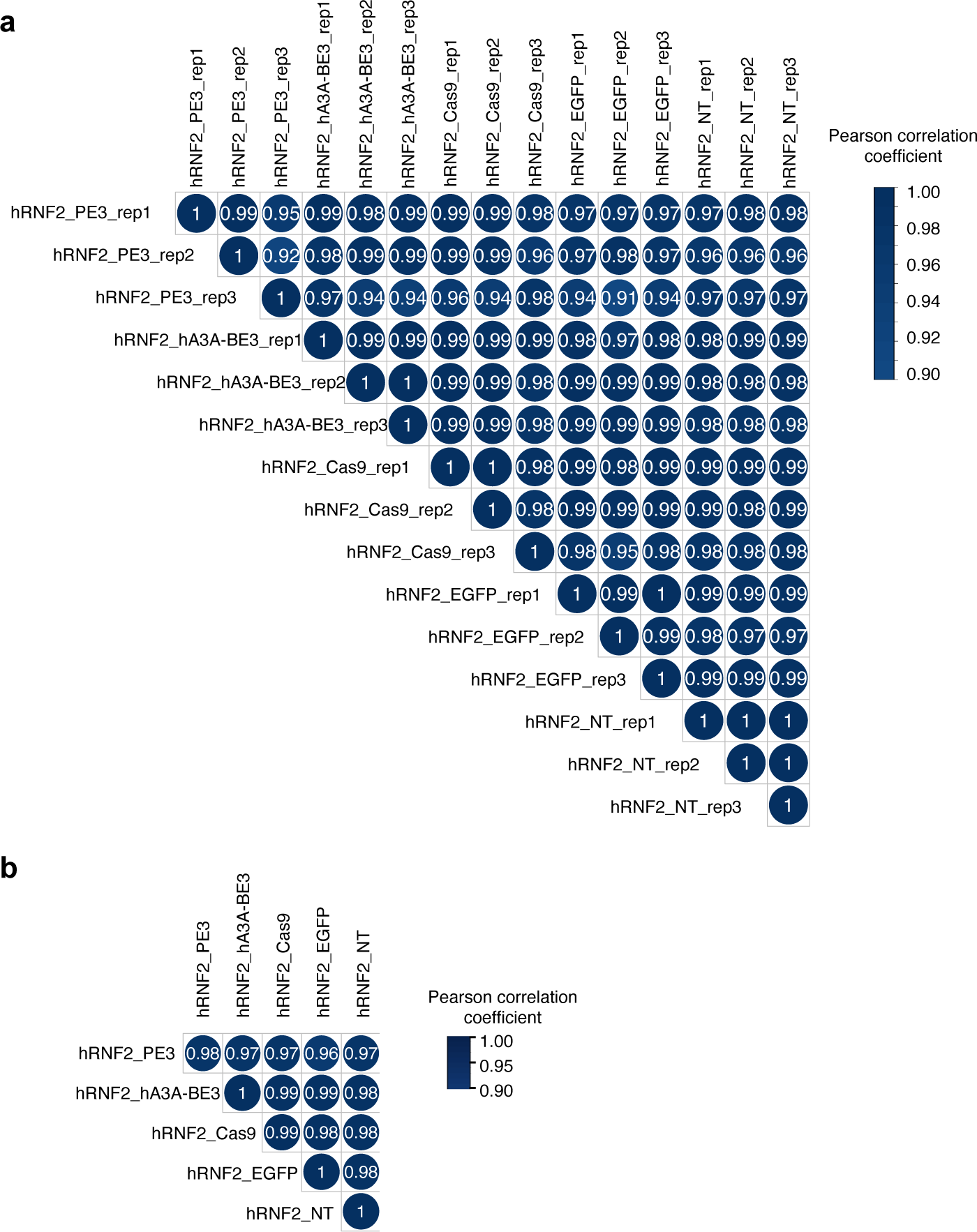
PE3 did not alter gene expression pattern in wild-type 293FT cells. (**a**) Heatmap of Pearson correlations on gene expression level (FPKM, fragments per kilobase of transcript per million mapped fragments) of WT 293FT cells treated with hA3A-BE3, PE3, Cas9, EGFP or left non-treatment (NT). (**b**) Heatmap of mean Pearson correlations coefficient across the same treatment on gene expression level (FPKM) of WT 293FT cells treated with hA3A-BE3, PE3, Cas9, EGFP or left nontreatment (NT).

## SUPPLEMENTARY TABLES

**Supplementary Table 1:**
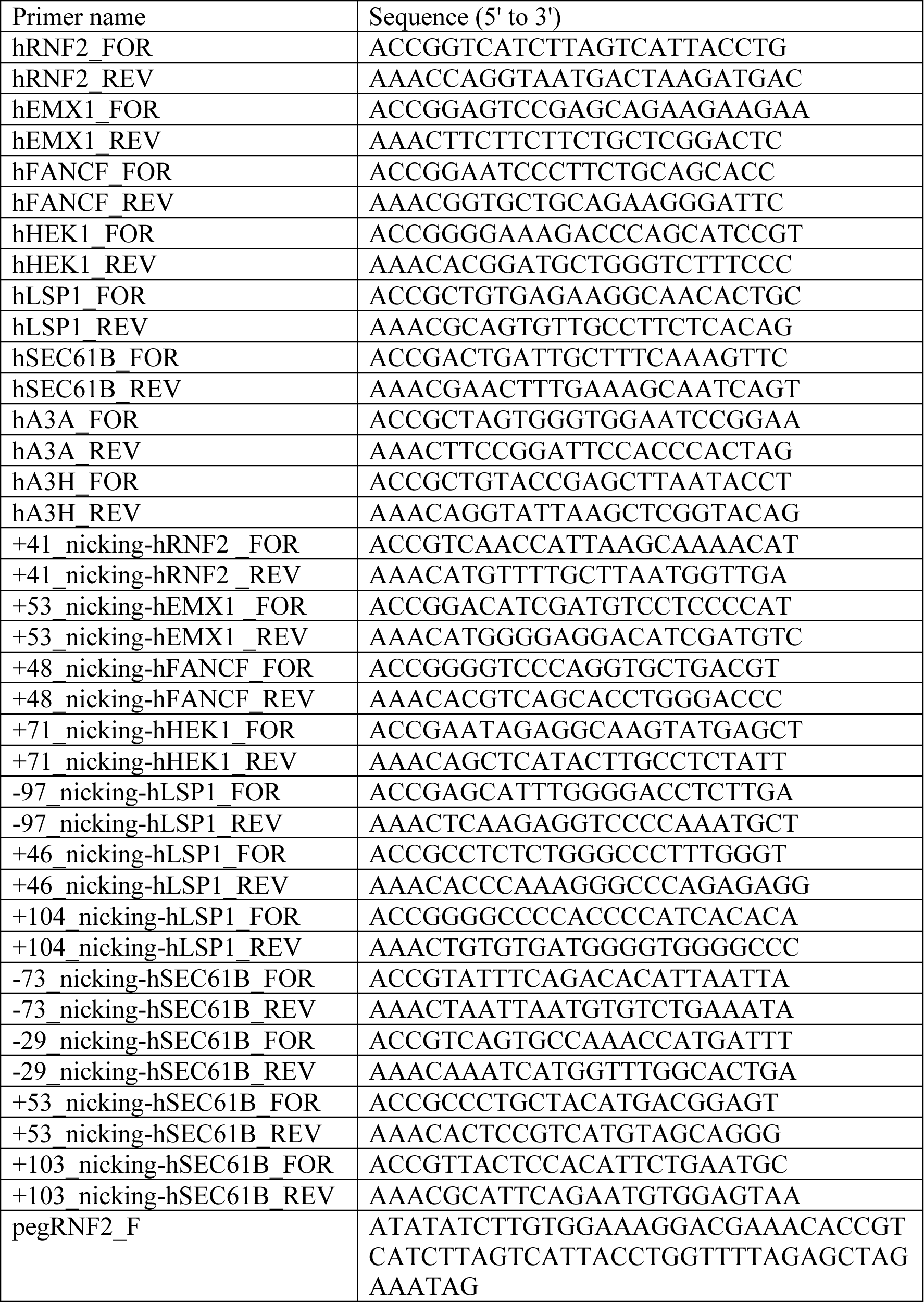
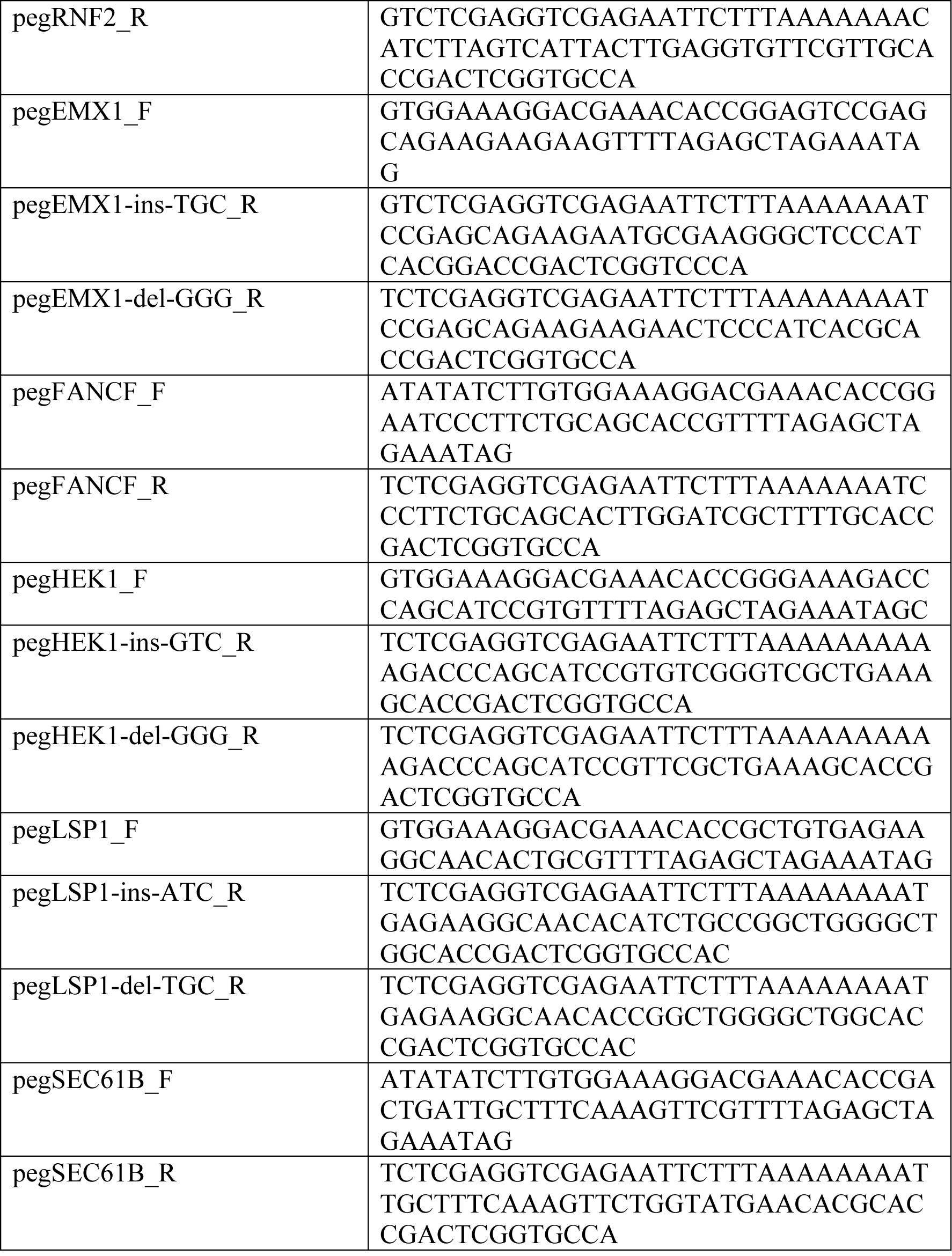
Oligos used for plasmid construction.

**Supplementary Table 2:**
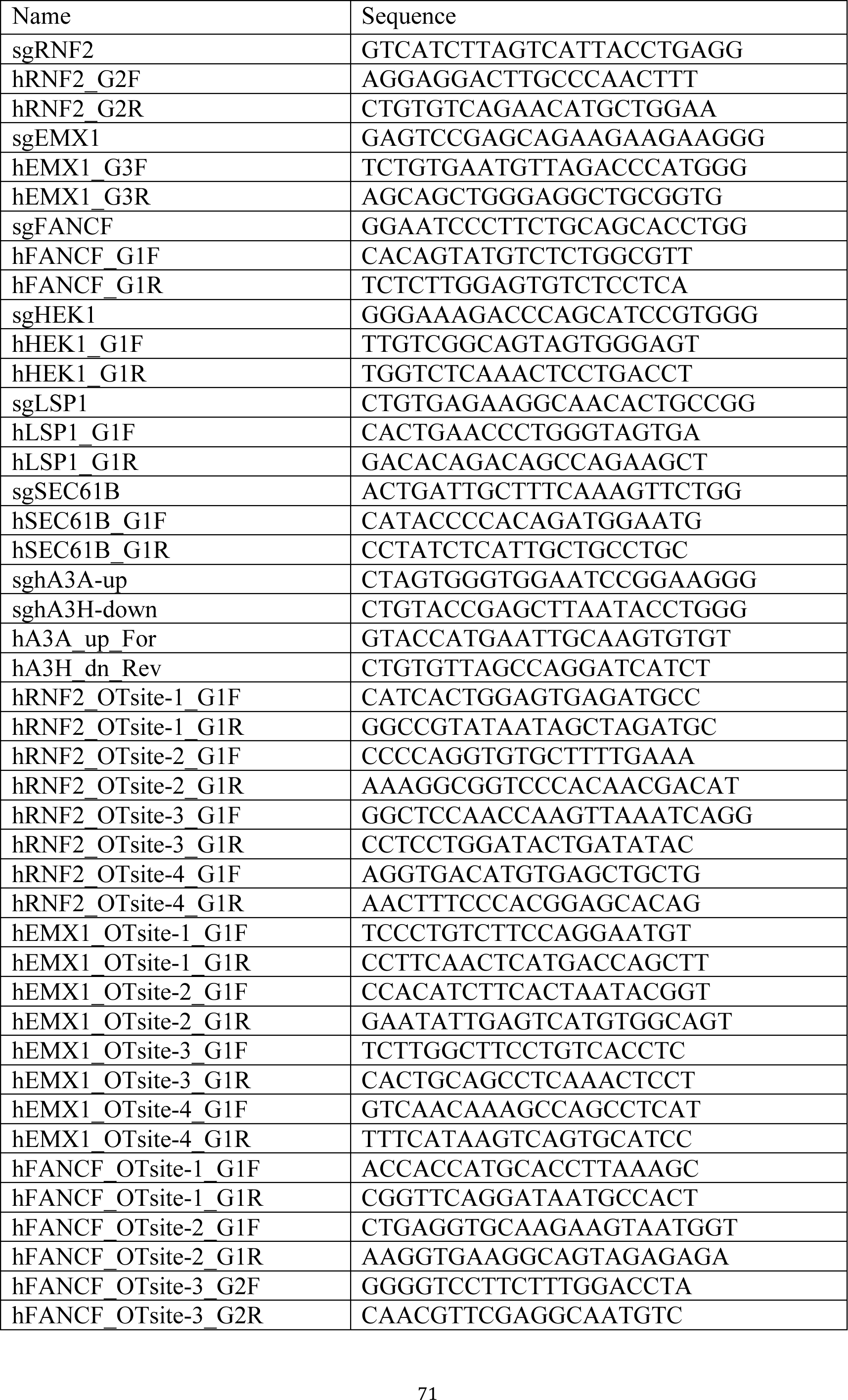
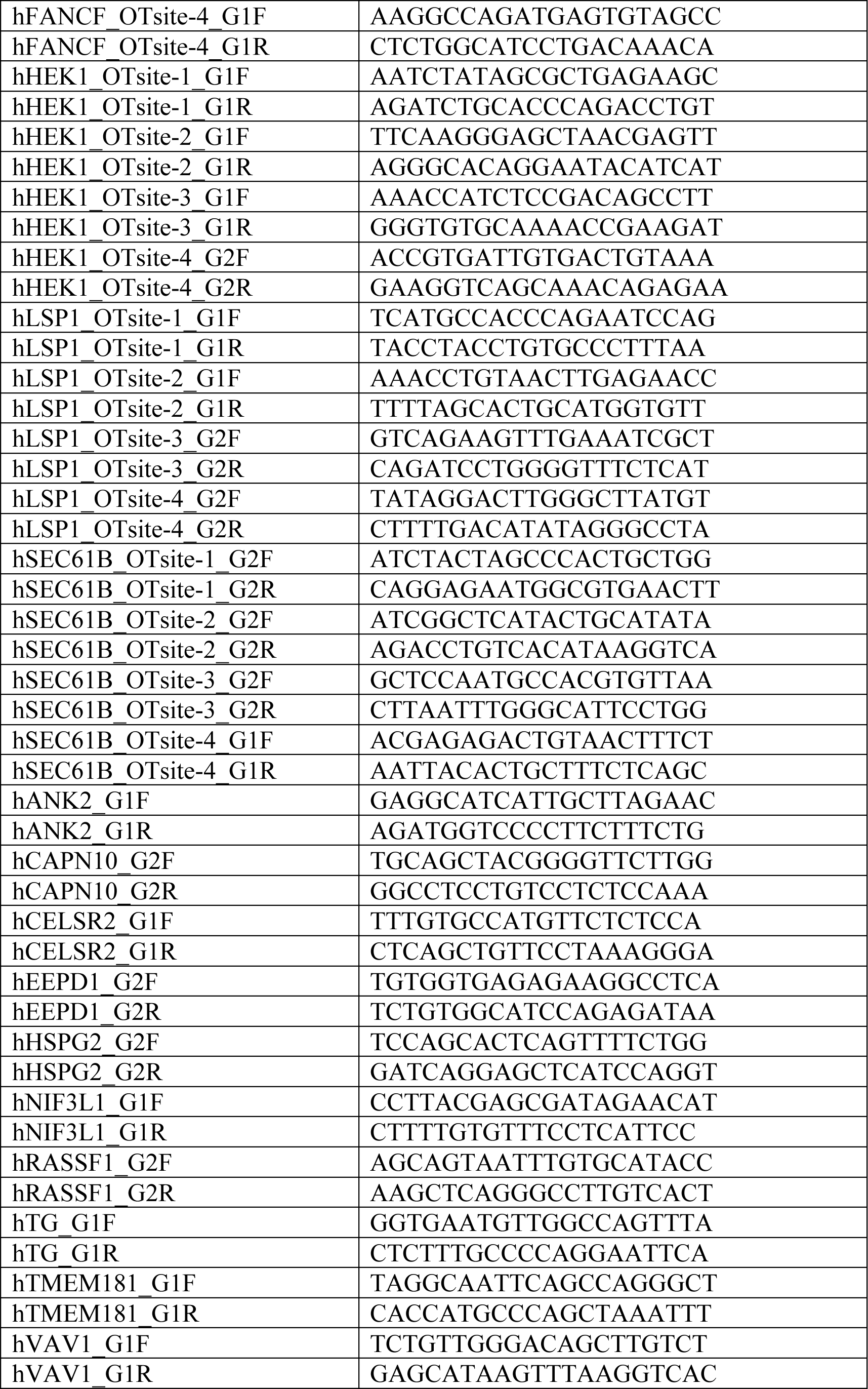
gRNA target sequences and PCR primer sequences for genomic DNA amplification.

**Supplementary Table 3: Calculation of on-target base substitutions.** Base substitutions were determined by deep sequencing and base substitution frequencies at the positions in gRNA-targeted genomic loci were calculated for indicated conditions. Read counts for all four types of bases are listed.

**Supplementary Table 4: Calculation of on-target indels.** Indels were determined by deep sequencing and indel frequencies at the examined gRNA-targeted genomic loci were calculated for indicated conditions. Counts of indel-containing reads and total mapped reads are listed.

**Supplementary Table 5: Calculation of gRNA-dependent OT base substitutions and indels.** Base substitutions and indels were determined by deep sequencing and total base substitution frequencies and indel frequencies at the examined gRNA-dependent OT genomic loci were calculated for indicated conditions. Read counts for all four types of bases and counts of indel-containing reads and total mapped reads are listed.

**Supplementary Table 6: Genome-wide base substitutions and indels.** Genomewide base substitutions and indels were determined by whole-genome sequencing for indicated single colonies.

**Supplementary Table 7: Calculation of telomere lengths and the numbers of telomeric repeat variants.** Telomere lengths and the numbers of telomeric repeat variants were determined by whole-genome sequencing for indicated single colonies.

**Supplementary Table 8: Transciptome-wide mutations.** Transcriptome-wide mutations were determined by whole-transcriptome sequencing for indicated conditions .

